# Gut immunity and the bacterial and eukaryotic microbiome of wild house mice

**DOI:** 10.64898/2026.04.21.719812

**Authors:** Louise Cheynel, Simon Hunter-Barnett, Lukasz Lukomski, Chris Law, Lorenzo Ressel, Jane Hurst, Mark Viney

## Abstract

It is well established that the microbiome can have major effects on animal biology. Here we have investigated the composition of the bacterial and eukaryotic gut microbiome of wild mice from three sample sites and sought to understand what affects its composition. We find that the bacterial and eukaryotic microbiome differs among mice from different sites. Among mouse traits, we found that only gut inflammation and the concentration of faecal immunoglobulin A affected the microbiome diversity. However, the microbiome diversity was more commonly affected by the microbiome composition itself, both within-bacterial and within-eukaryotic, but also by cross bacterial-eukaryotic effects. We found that most hosts produce IgA that binds some of their gut bacteria, though mice are largely idiosyncratic in which taxa they bind with IgA, with a few taxa commonly IgA-bound. The eukaryotic microbiome was dominated by fungal taxa, and included *Eimeria* infection that was particularly common at one of the sites. At the high *Eimeria* prevalence site, mice had comparatively marked caecal inflammation and significantly greater IgA responses. Our results emphasise the substantial among-individual mouse differences in gut microbiome composition, gut physiology and immunology, and the biological significance of the bacterial-eukaryotic effects that we suggest requires further study.

## 1. Introduction

It is now well established that the gut microbiome can have profound effects on its host. Most of the work discovering this has come from the study of laboratory animals, livestock and humans. There are relatively few studies of wild animals and their microbiomes, yet the expectation must be that their biology is also significantly affected by their microbiomes.

There have some studies of the microbiome of wild mice, though far fewer than of laboratory mice. A comprehensive comparison of wild and laboratory house mice (*Mus musculus*) has shown that their gut microbiomes are quite different, with those of the wild mice generally more diverse, and with greater among-individual variation, than in laboratory mice [1]. These findings are consistent with other studies that moved wild house mice to the laboratory, finding that the microbiomes diverged with this transfer [2]. Other wild-to-laboratory transfers have shown that wild house mouse populations differ in their microbiomes; on transfer to the laboratory there is a reduction in microbiome diversity, but mouse population differences persist, suggesting a host genetic basis [3]. Analogous observations of wild and laboratory deer mice (*Peromyscus maniculatus*) also found greater microbiome diversity in wild mice than in laboratory-maintained animals, and evidence of comparatively greater temporal turnover in wild mice [4]. Studies in wood mice (*Apodemus sylvaticus*) found relatively subtle differences between the microbiome of a laboratory-established population and wild populations, but differences among the wild populations [5]. Similarly, in bank voles (*Clethrionomys glareolus*), urban-to-rural transfer showed that the microbiome composition of the microbiome was affected by both the original and new site of the voles [6]. A complementary approach to these studies is to transplant wild mouse microbiomes into laboratory mice, so-called wilding mice [7]. These transplants recapitulate a number of aspects of wild mouse biology and immunology in the transplanted mice [8]. Rewilding of laboratory mice into outdoor enclosures also changes the gut microbiome, making it more diverse and increasing the fungal component [9–10].

Factors other than the wild *vs*. laboratory location of mice can affect the microbiome. For example, in wood mice (*A. sylvaticus*), social contact between animals, their shared use of space, similarity of habitats, and their age all contributed to gut microbiome similarity [11]. Other studies of wild *A. sylvaticus* found evidence for seasonal effects on the microbiome, on top of otherwise idiosyncratic microbiomes [5]. Also in *A. sylvaticus*, early life maternal–offspring contact increased the microbiome diversity and richness [12].

While acquisition of microbial taxa from the environment is a necessary first step for incorporation into the microbiome, the persistence and relative abundance of specific taxa will then depend on within-host processes. These processes will include the effects of host physiology (and particularly gut physiology for the gut microbiome), the host immune response, as well as interactions among microbial taxa themselves. Extensive studies, particularly in laboratory mice, have shown how various host-factors can affect component taxa of the microbiome, but there has been very little study of this in wild animals.

Many specific mechanisms affecting the microbiome can be studied in laboratory systems, allowing precise dissection of the effects of components of host biology on the gut microbiome. While this is a powerful way to investigate biological mechanisms, it is now well-established that there are considerable differences between wild and laboratory animals, particularly in the immune responses they make [8,13–15]. Immunologically, studies of wild animals have shown that they have humoral and cellular immune responses consistent with extensive antigenic exposure, and exposure that is greater than laboratory animals’, and that wild mice are genetically diverse in genes whose products have immunological function [16–17]. However, wild animals’ *in vitro* proliferative responses are typically depressed compared with those of laboratory animals. This may be to avoid immunopathology that might otherwise occur given their very high antigenic exposure. A consistent theme of a body of work using rewilded and wilding mice is that a wild mouse microbiome results in an experienced immune system, which contrasts with the comparatively naïve, antigen-inexperienced immune system of conventional laboratory mice [8,18–19]. The increase in the fungal component of the gut microbiome of rewilded mice drives particular change in the rewilded animals’ immune state [10].

The host immune response, particularly humoral responses, may directly affect components of the gut microbiome. In the mammalian gut and at other mucosal surfaces, large quantities of IgA is secreted (*c*.5 g per day into the human gut), which acts as a mucosal site barrier. Work with laboratory mice has shown that IgA binds to a sub-set of the bacterial microbiome and that this can affect those bacteria [20–21]. Bacterial taxa bound by IgA are thought to be those that are pathogenic, or potentially pathogenic, to their host with the IgA response made to ameliorate that threat. But what microbiome taxa are targeted by IgA in wild mice has not yet been investigated. However, wild house mice can have very substantially higher gut [IgA] than laboratory animals [14], suggesting that the IgA binding of bacteria may be comparatively more pronounced in wild animals. Further, it has previously been suggested that such IgA responses may act as a selective force against IgA bacteria, driving the mode and tempo of their evolution [21].

The principal focus of gut microbiome research is the bacterial microbiome. However, eukaryotic taxa are a substantial proportion of the gut microbiome, but have received very much less attention [22]. Among wild animals there are a number of eukaryotic pathogens that can cause significant morbidity and mortality. There is also the potential for interactions between eukaryotic and bacterial components of the microbiome to affect the composition and diversity of the microbiome. Indeed, it is of note that rewilded and wilding mice have a greater fungal component of their microbiome compared with laboratory mice [10–18]. Studies in laboratory mice have also shown how some fungal taxa can have very significant effects on immune responses [23]. The greater fungal component of the microbiome in wild, rewilded or wildling mice compared with laboratory mice might allow greater interactions between the bacterial and eukaryotic microbiome either directly or be mediated by host physiology and immunology.

The overall aim of this work was to investigate the composition of the bacterial and eukaryotic microbiome of wild mice and to understand how its composition is affected by aspects of animals’ biology. We specifically aimed to determine: (i) the composition of both the bacterial and eukaryotic gut microbiome; (ii) the immune state of the gut of wild animals and how, if at all, this affects the composition of the microbiome; (iii) what components of the bacterial microbiome is bound by IgA; (iv) if the bacterial and eukaryotic components of the microbiome interact, and if so to what effect.

## 2. Material and methods

### 2.1. Mice

We live trapped 58 mice (*Mus musculus domesticus*) from 3 sites in England between July and November 2021; specifically, a pig farm in Nottinghamshire; horse stables near Southport, Merseyside; and a dairy farm on the Wirral Peninsula (**Supplementary Information 1**).

Mice were killed, sexed, weighed, and measured from the tip of their snout to the base of their tail. We calculated Body Mass Index (BMI) as log body mass /log body length, which predicts fat mass of mice [24]. We aged mice as [25]. Faecal samples were collected both from the trap in which a mouse was caught and directly from the dissected terminal portion on the gut. The caecum was separated from the small and large intestine, opened along the greater curvature [26], and the contents washed into 2 mL of PBS. Faecal samples and caecal contents were stored at −80°C no more than 2 hours after the mouse was killed. We used faecal samples for analysis of bacteria by 16S rRNA sequencing; we used faecal and caecal samples for analysis of eukaryotes by 18S rRNA sequencing. All intestinal tissues were fixed in 10 % v/v formal saline after dissection. The distribution of individuals by study site, sex, and age is in **Supplementary Information 1**. This work was reviewed and approved by the University of Liverpool Animal Welfare and Ethical Review Board.

### 2.2 Intestinal antibody response

We measured the faecal concentration of total immunoglobulin A (IgA) as described in **Supplementary Information 2**.

### 2.3. Faecal mucin assay

Mucins are a component of the mammalian mucosal protection [27] that can be quantified by the faecal concentration of N-acetylgalactosamine (GalNAc) [28]. GalNAc concentration was measured using a commercially available fluorometric assay kit (Cosmo Bio; [29]), following the manufacturer’s methods [30], with three mouse controls: (i) a male specific-pathogen free (SPF) C57BL/6, (ii) a female non-SPF C57BL/6, and (iii) a female from a captive colony derived from wild house mice.

### 2.4. Assessment of gut inflammation

We measured gut inflammation histologically in five sections of gut tissue for each mouse and for control, laboratory mice (i – iii, above): 1cm of the small intestine (one replicate per mouse); 0.5cm of the large intestine (two replicates per mouse); 0.2cm of the caecum (two replicates per mouse), which were used to produce longitudinal sections of the large and small intestines and transverse sections of the caecum, from which haematoxylin and eosin stained sections were produced. There were examined for evidence of inflammation after [31], which is a combined measure of the severity and extent of immune cell infiltrate in the lamina propria; hyperplasia and goblet cell loss; ulceration and crypt loss; and, for the small intestine, villous blunting and/or atrophy following exposure of luminal antigens [30]. For the two replicates of the caecum and of the large intestine, the larger of the two values was used. Some samples failed to process so that the final sample sizes for 58 wild mice were 57, 55, 53 for the small intestine, caecum, and large intestine, respectively.

### 2.5. *Eimeria* infection

*Eimeria* is a common parasite of wild rodents [32–33] and we determined its presence in the mice by PCR assay of faecal and caecal DNA, as described in **Supplementary Information 2**.

### 2.6. Flow cytometry counting and sorting of IgA^+^ and IgA^−^ faecal bacteria

We used flow cytometry to count the bacterial load and to collect three bacterial fractions: (i) the pre-sort, (ii) bacteria coated with IgA (henceforth IgA^+^) and (iii) those not (IgA^−^), by combining and adapting previous methods [20,34] and from which DNA was extracted, all as described in **Supplementary Information 2**.

### 2.7. 16S rRNA sequencing and bioinformatic analyses

We amplified the V4 region of the 16S rRNA coding sequence and sequenced these on an Illumina miSeq using barcoded primers (primer pair F515/R806; [35]. We used a ZymoBIOMICS Microbial Community DNA Standard (Zymo, D6305) mock community control. Sequences were processed using the DADA2 pipeline in R as described in **Supplementary Information 2**, resulting in 2,383 Amplicon Sequence Variants (ASVs).

### 2.8. 18S rRNA sequencing and bioinformatic analyses

We characterised the eukaryotic component of the caecal and faecal microbiome by 18S rRNA sequencing (primers 528F and 707R;Novogene, 2023) using faecal and caecal DNA. The sequenced amplicons generated 10,477,596 reads for all samples. Controls were: (i) negative, consisting of DNA extraction of a blank sample [36], and (ii) positive, consisting of a *Saccharomyces cerevisiae* and *Cryptococcus neoformans* (ZymoBIOMICS) microbial community.

We identified taxonomically the sequence reads as described in **Supplementary Information 2**. ASVs were agglomerated at the species level to simplify analyses, retaining any unassigned taxa (phyloseq package). ASVs that occurred fewer than ten times across the entire dataset were removed, following [37]. ASVs assigned to Phragmoplastophyta, Chlorophyta, Klebsormidiophyceae (plants); Arthropoda (insects); and Vertebrata (host) were removed. Our negative control identified a single ASV, present in 17 of the wild mouse samples; this ASV was excluded from all further analyses. We successfully recovered the species present in the mock communities.

### 2.9. Statistical analyses

To investigate the covariation among measured traits, we performed a principal component analysis (PCA) of 17 intestinal variables; specifically (i) [faecal IgA], (ii) the proportion of bacteria that were IgA^+^, (iii) total bacterial load, (iv-vi) inflammation of the small intestine, large intestine and caecum, (vii) [faecal mucin], (viii-xv) relative abundance of Ascomycota, Basidiomycota, Apicomplexa and Nematoda in the caecum and in the faeces, (xvi) BMI, (xvii) age. A total of eight mice were excluded because they had some data missing for at least one of the 17 variables. The PCA was performed using the R package “ade4” [38].

To determine if an ASV was preferentially in the IgA^+^ or IgA^−^ fraction we calculated the IgA probability ratio scores for each ASV after [39]; if an ASV was absent from a mouse, then its probability is zero and so we excluded all zero IgA probabilities. We focussed on the 500 most abundant ASVs, which together accounted for *c*.90% of the total relative abundance. We used the number of sequence reads within each ASV as a measure of the abundance of that ASV.

We calculated bacterial and eukaryotic microbiome alpha diversity (*i.e*. diversity within an individual) using Shannon’s index after [40]. For the bacterial microbiome we compared alpha-diversity in the pre-sort, IgA^+^, and IgA^−^ fractions and between sites using Wilcoxon signed-rank tests. To investigate what factors affected the alpha diversity we constructed step-wise linear models, retaining only significant factors at each step. We first tested (i) mouse traits (site, sex, age, BMI, reproductive status), (ii) immune related traits ([faecal IgA], bacterial load, proportion IgA^+^ bacteria), (iii) gut inflammation ([mucin], small intestine, large intestine and caecum inflammation), (iv) the faecal eukaryotic microbiome (relative abundance of Apicomplexa, Basidiomycota, Nematoda, Ascomycota), (v) the caecal eukaryotic microbiome (relative abundance of Apicomplexa, Basidiomycota, Nematoda, Ascomycota), and (vi) the main bacterial phyla (relative abundance of Firmicutes, Bacteroidota, Actinobacteriota and Proteobacteria). At each step, we used a model selection procedure based on the Akaike Information Criterion (AIC) [41]. We retained the model with the lowest AIC, and when the difference of AICs between competing models was less than 2, we retained the model with the fewest parameters to satisfy parsimony rules [41]. Finally, the goodness of fit of the selected models was assessed through calculating conditional (*i.e*. total variance explained by the best supported model) R^2^ formulations.

For beta diversify we calculated pairwise dissimilarities among samples using Bray-Curtis [42–43]. For the bacterial microbiome beta diversity we normalized read abundance to compositional proportion data and performed a principal coordinate analysis (PCoA) of the Bray-Curtis distances, to visualize compositional differences between sites. We then conducted permutational multi-variate analysis of variance (PERMANOVA, with 9999 permutations) tests on beta diversity for the effects of the same factors tested for alpha diversity (above), but also testing for the effects of bacterial and eukaryotic alpha diversity, and of the abundance of the 12 ASVs that are preferentially IgA^+^. For both the analyses of alpha and beta diversity we ran separate analyses for the bacterial (pre-sort) microbiome, the caecal eukaryotic microbiome, and the faecal eukaryotic microbiome.

We constructed networks of the 75 most abundant ASVs from the bacterial (pre-sport) and eukaryotic microbiome using NetCoMi R [44] and then SPRING [45] to estimate associations between taxa. We constructed a network for the whole data set and then separately for data from each of the 3 sample sites.

To compare the *Eimeria* infection status of faecal and caecal samples we used Pearson’s and McNemar Chi-squared tests. To compare the number of *Eimeria-* infected mice among sites we used Fisher’s exact test with subsequent pair-wise tests, accounting for multiple testing after [46], where a *p* < 0.05 indicates a difference in the proportion of *Eimeria-*infected mice among the sampling sites. To compare the [faecal IgA], [faecal mucin], and intestinal inflammation between *Eimeria*-positive and *Eimeria*-negative mice we used a Wilcoxon two-sample test, a generalised mixed-effects model (GLMM, [47]), and a Fisher’s exact test, respectively. The GLMM incorporated the assay plate number, faecal starting weight and faecal source (trap or intestine) as random effect terms, and used a Gamma error distribution.

## 3. Results

### 3.1 Covariation among gut traits

We used a PCA to examine patterns of covariation among traits likely relevant to the composition of the gut microbiome of wild mice (**Figure 1; Supplementary Information 3**).

**Figure 1.**
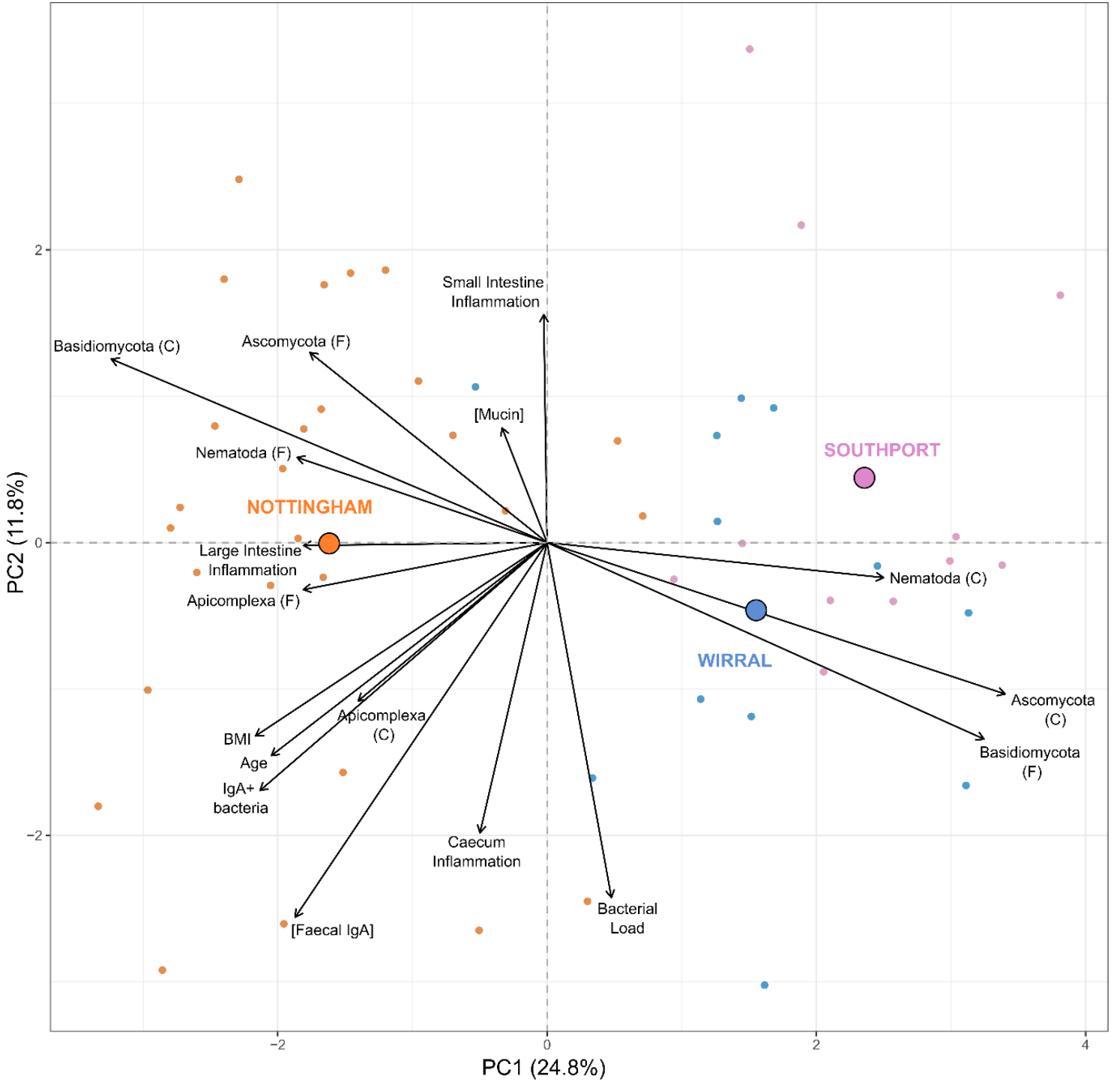
Principal component analysis. The first two principal components of a PCA analysis of 17 traits in mice from three sites (Nottingham, Southport and Wirral). Arrows indicate the traits’ contribution to each of the first two PCs, with longer arrows denoting stronger correlations. Small points are individual mice, and large points are average values for mice at each site, with each site and mice from that site colour coded. PC1 accounts for 25% of the variance in the data, and PCs 2-4 each for a further 10% (**Supplementary Information 3 to 4**).

Mouse age and BMI both strongly correlate with PCs 1-3, and age and BMI are strongly positively correlated (**Figure 1; Supplementary Information 3**). The [faecal IgA] also strongly correlates with PCs1 and 2, and is positively correlated with mouse age and BMI (**Supplementary Information 4**). Other aspects of the microbiome are also correlated with PC1, specifically the proportion of IgA^+^ bacteria, the abundance of Ascomycota (caecal), Basidiomycota (caecal and faecal) and Nematoda (faecal) (**Supplementary Information 3**).

For nematodes, Ascomycota and Basidiomycota each of their caecal and faecal relative abundances are differently loaded onto PCs1 and 2, and for each taxonomic group the caecal and faecal measures are negatively correlated (**Supplementary Information 3**). In contrast the caecal and faecal measures of Apicomplexa (which is principally *Eimeria*, below) is more similarly loaded onto the PCs, and is positively correlated (**Figure 1; Supplementary Information 3**).

The measures of inflammation of each of the three gut regions are distributed across the PCs and they correlate poorly (**Supplementary Information 3**). The [faecal mucin] is similarly distributed across the PCs, though it does correlate with large intestine inflammation. There is correlation between measures of Apicomplexa infection and gut inflammation; specifically Apicomplexa (caecal and faecal) and large intestine inflammation; Apicomplexa (faecal) and caecal inflammation. Basidiomycota (caecal) also correlates with widespread inflammation, in the small and large intestine and in the caecum.

Concerning how mice vary, they are generally distributed across PCs 1 and 2 by the site they originated from (mean ± SD PC1 and PC2 respectively for: Nottingham −1.64 ± 1.08; 0.02 ± 1.51; Southport 2.34 ± 0.89; 0.46 ± 1.33; Wirral 1.53 ± 1.04, −0.45 ± 1.29), and there are no outliers. PC1 principally separates Nottingham mice from Southport and Wirral mice (**Figure 1**). There is no major separation of young *vs*. adult (< 6 *vs*. ≥ 6 weeks old; 32 and 25 respectively) mice, or male *vs*. female mice (**Supplementary Information 3)**.

### 3.2 The bacterial microbiome

The bacterial microbiome is dominated by bacteria of the Bacteroidota and Firmicutes phyla, but this varies substantially among mice **(Figure 2)**. The proportion of Bacteroidota varies from 3-87% (mean 35%), as too the proportion of Firmicutes 9-90% (mean 57%). Bacteria belonging to the phylum Actinobacteriota occur in most mice, with an average relative abundance of 2.5% (range 0.1-22%). Proteobacteria were more abundant in Southport and Wirral populations (2.6 ± 4.4%; 2.2 ± 2.9% mean ± SD, respectively) than in Nottingham (1.3 ± 1.5%); *vice versa* Actinobacteriota are generally more abundant in Nottingham-derived mice (3.3 ± 5.1%) than in those from Southport and the Wirral (1.8 ± 2.4%; 1.2 ± 0.75%, respectively). This among sample site variation in the abundance of different bacteria taxa is consistent with the among sample site variation in gut traits (above).

**Figure 2.**
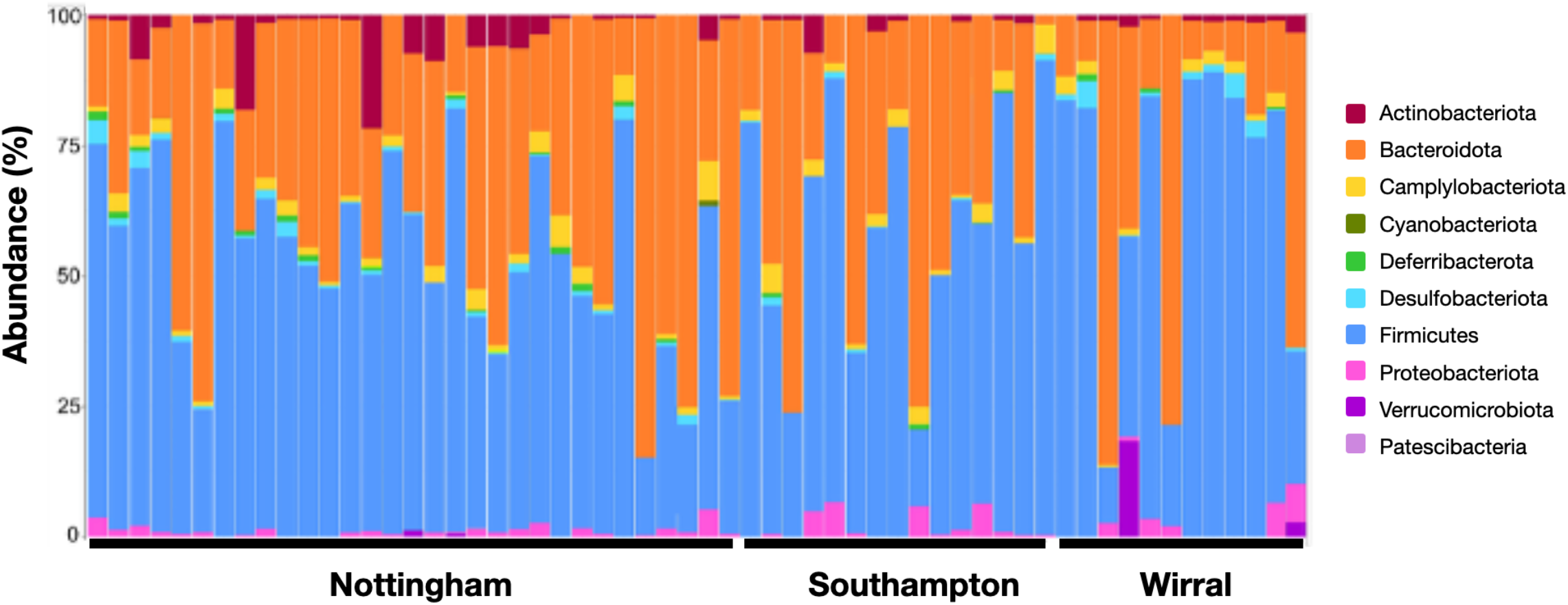
Bacterial microbiome. The relative abundance of taxa from each phylum for wild mice from 3 sites, (Nottingham, Southport and Wirral).

In all we resolved 2,383 ASVs, though 90% of the total relative abundance is in the 500 most abundant ASVs. The mice differed in the number and combination of ASVs that they harboured, with an average of 168 ASVs (SD 47; range 54-263) harboured by each mouse. There was variation in the bacterial load of each mouse ranging from 6×10^6^ to 3×10^8^ per gramme of faeces (6 ± 4 x 10^7^, mean ± SD), and a positive relationship between bacteria load and [faecal IgA] (**Supplementary Information 4**).

The bacterial microbiome alpha diversity did not differ among mice from the three sites (Nottingham 4.20 ± 0.62; Southport 4.00 ± 0.48; Wirral 3.97 ± 0.59, mean ± SD; Kruskal-Wallis: χ^2^_2_ = 3.01, *p* > 0.05) (**Supplementary Information 5**). In contrast, 13% of the beta diversity variation was due to sample site (PERMANOVA: F2 = 4.19, R2 = 0.132, *p* = 0.001; **Supplementary Information 6**), and it differed between the sites (p < 0.01 for the three pairwise comparisons).

We investigated what factors affected the alpha diversity of the bacterial microbiome, by constructing stepwise linear models, retaining significant factors at each step, testing in sequence for: (i) mouse traits, (ii) immune-related traits, (iii) gut inflammation, (iv) the faecal eukaryotic microbiome, (v) the caecal eukaryotic microbiome, and (vi) the main bacterial phyla. We found only a significant negative effect of the main bacterial phyla, specifically the relative abundance of Bacteroidota, on alpha diversity (**Supplementary information 7**).

We used PERMANOVA to investigate factors that affected the beta diversity, and identified 9 factors that each accounted for ≥5% of the total variation in beta diversity, specifically: (a) two mouse traits – sample site and age (< 6 weeks old *vs*. ≥ 6 weeks old); (b) three bacterial microbiome traits – relative abundance in the pre-sort fraction of Firmicutes and of Bacteroidota, bacterial microbiome alpha diversity; and (c) four eukaryotic microbiome traits – relative abundance of caecal Ascomycota, caecal Basidiomycota, faecal Basidiomycota, caecal Nematode (**Supplementary Information 8**). We especially note that the beta diversity of the bacterial microbiome is affected by the faecal and caecal eukaryotic microbiome.

### 3.3 IgA binding to the bacterial microbiome

Using flow cytometry we determined that an average of 16% (SD 6.4%; range 4-29%) of a mouse’s bacterial microbiome was IgA^+^ (**Figure 3**). The alpha diversity did not differ between the IgA^+^, IgA^−^ and pre-sort fractions (IgA^+^ 3.87 ± 0.67; IgA^−^ 3.94 ± 0.65, pre-sort 4.10 ± 0.58, mean ± SD) (**Supplementary Information 9**). The beta diversity of the pre-sort, IgA^+^ and IgA^−^ fractions were generally not dissimilar from each other for each mouse, suggesting that each mouse tends to have an individual signature in its bacterial microbiome beta diversity (**Supplementary Information 10**).

**Figure 3.**
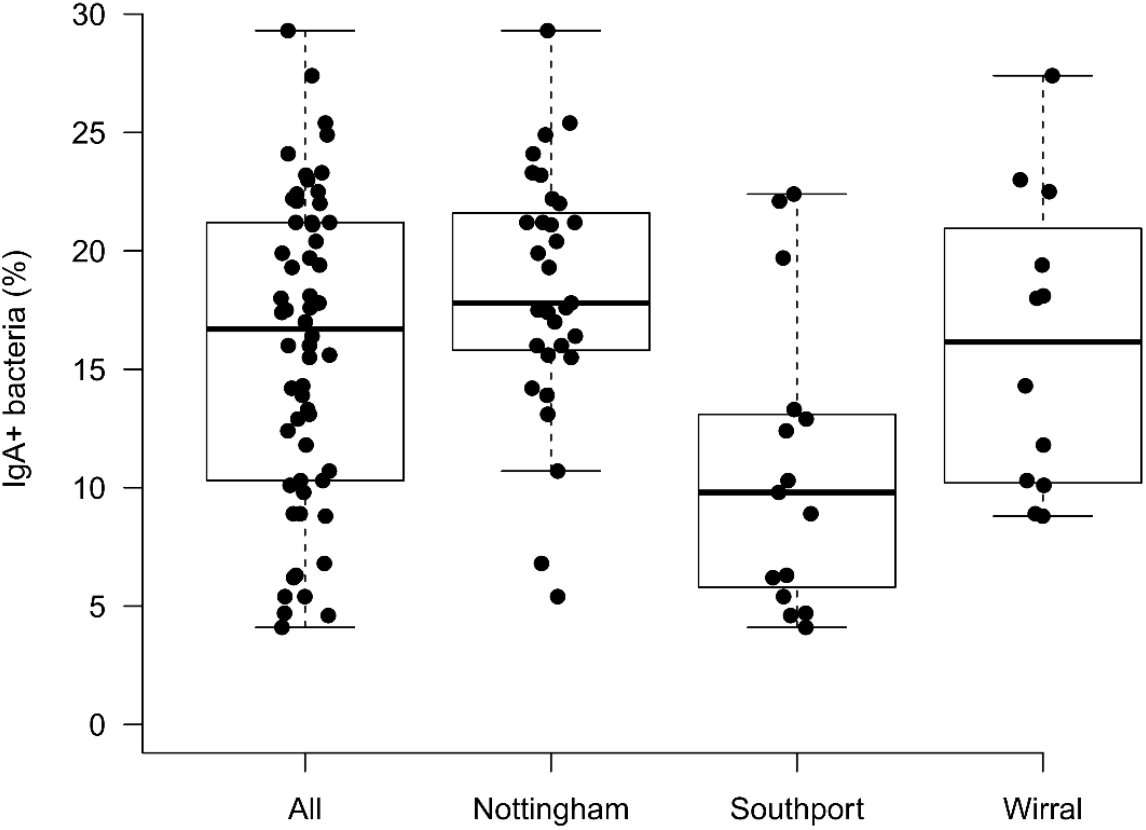
IgA^+^ bacteria. The proportion of bacteria that were bound by IgA (IgA^+^) for wild mice from 3 sites. The horizontal line is the median, the box the inter-quartile range and the whiskers show the minimum to maximum values in the dataset.

We calculated the IgA probabilities for each mouse for the ASVs that it harboured, finding that most IgA probabilities were negative (88%), with just 12% positive (**Figure 4A**). Mice were not consistent in binding IgA to an ASV, with no ASV having a positive IgA probability in all mice (**Supplementary Information 11**).

**Figure 4.**
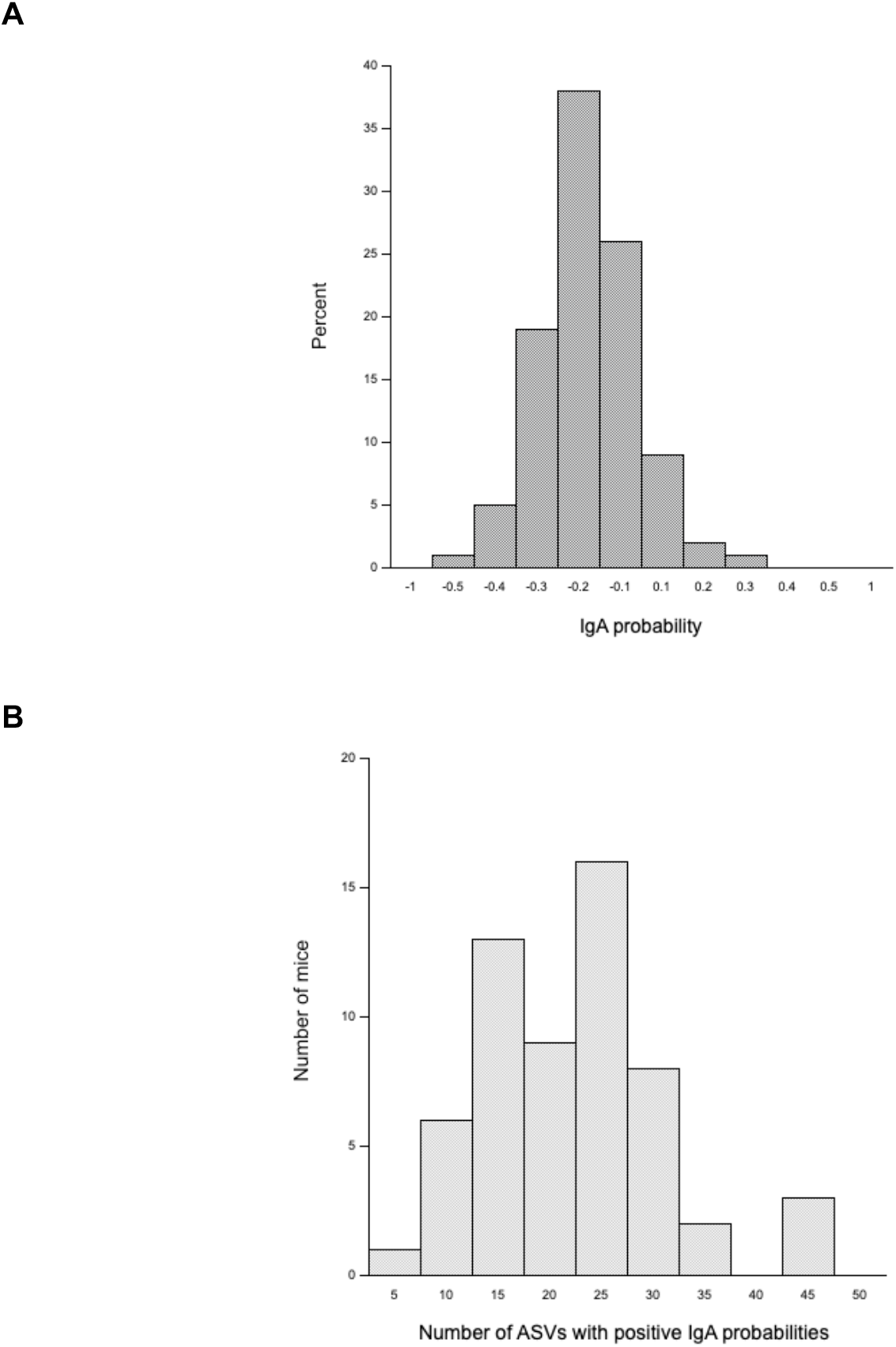
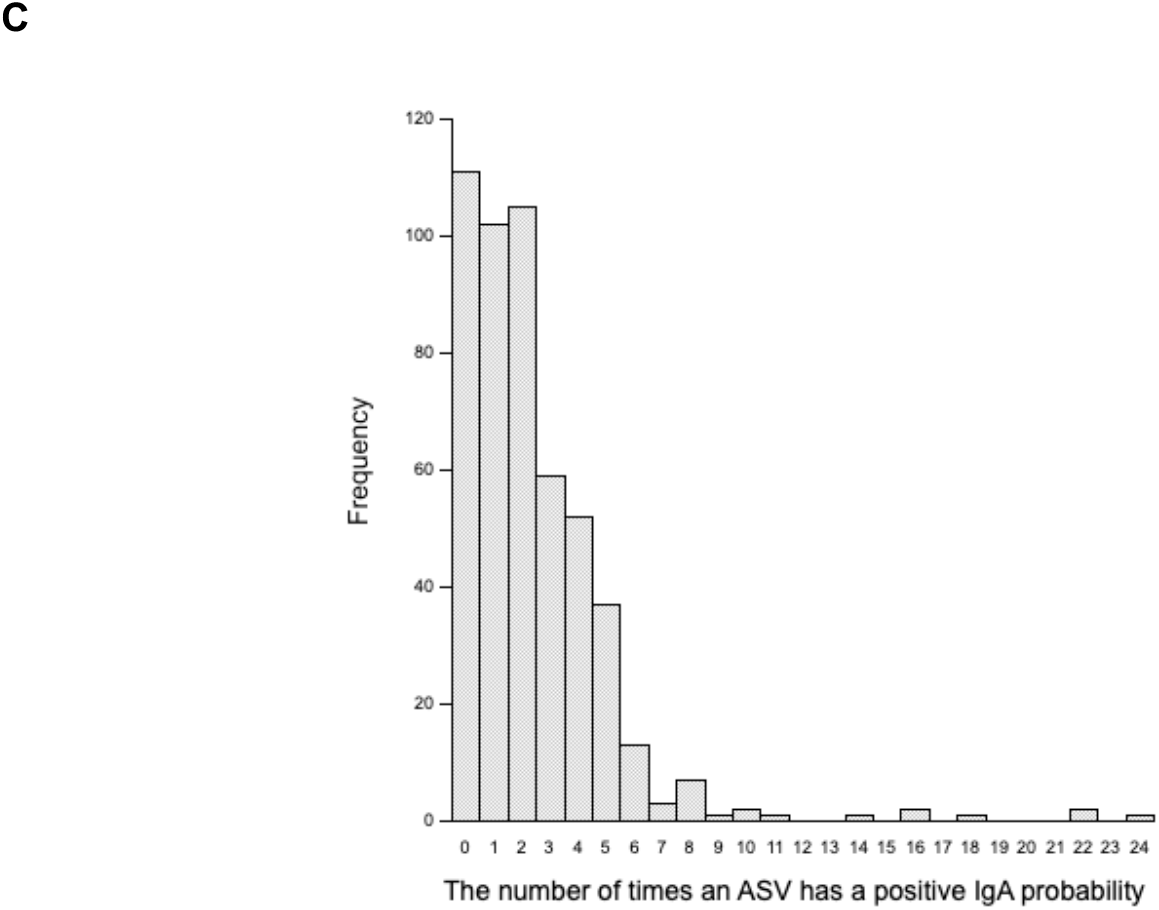
IgA probabilities. (A) The distribution of IgA probabilities from 58 mice for the 500 most abundant ASVs; the x-axis scale is in 0.1 increments, until ± 0.5 and then a final 0.5-sized increment. (B) The distribution of the number of mice and how many ASVs they harbour that have a positive IgA probability. (C) The distribution of the frequency with which ASVs have a positive IgA probability in a mouse.

We asked how mice varied in the number of ASVs that they harboured that had a positive IgA probability, finding that 19 mice (34%) account for 49% of all ASVs with positive IgA probabilities (**Figure 4B**). We identified these 19 mice on the PCA of traits likely to be relevant to the composition of the gut microbiome, finding that these 19 mice were widely dispersed across the PCs 1 and 2 (**Supplementary Information 12**).

Mindful of the PCA-revealed relationships among mouse age, BMI, [faecal IgA], and proportion of bacteria that are IgA^+^, we asked if the number of positive IgA probability ASVs that any mouse harboured was related to these other factors. We found a positive relationship (R = 0.57) between the percent of bacteria that were IgA^+^ in a mouse and the number of positive IgA probability ASVs that a mouse harbours (**Supplementary Information 13**), but no relationship with mouse age, BMI, [faecal IgA], nor *Eimeria* abundance (Section 3.6, below).

We also asked how ASVs varied in their IgA probability, finding that a small number of ASVs more often had a positive IgA probability (**Figure 4C**). Specifically, 389 ASVs had a positive IgA probability at least once, and 34 ASVs had a positive IgA probability in at least 6 mice (which is *c*.10% of the sampled mice). We assessed the ASVs that most often had a positive IgA probability in two ways. First, as the 10 ASVs that most commonly had a positive IgA probability in a mouse. Second, as the sum of the IgA probabilities for each ASV across all mice, finding that 10 ASVs had such a value that was greater than the mean value + 2 standard deviations of this value. The ASVs (and their taxonomic classification) identified by these two criteria are shown in **Table 1**. This revealed a large overlap among ASVs identified in these two ways, with 8 ASVs matching both criteria, out of 12 in total. Further, this shows that these positive IgA probabilities are specific to ASVs, and do not overlap with other con-generic ASVs (**Table 1**). Despite these 12 ASVs commonly being harboured with a positive IgA probability, many other mice also harboured these ASVs but with a negative IgA probability. Specifically, for each of these 12 ASVs among the 58 mice, an average of 15 (range 8-24) mice had the ASV with a positive IgA probability, while an average of 12 (range 1-30) mice harboured the ASV with a negative IgA probability. There were many positive correlations of abundance of these 12 ASVs in the IgA^+^ fraction (23 ≥ 0.2), and more so than among 12 other randomly selected ASVs (4 > 0.2; **Supplementary Information 14**). There was no effect of the relative abundance of each of these 12 ASVs in the IgA^+^ fraction on the alpha or beta diversity of the bacterial microbiome.

**Table 1.** Bacterial ASVs that are preferentially bound by IgA. In ‘Common’ the shading shows the 10 ASVs that most often have a positive IgA probability, with the number indicating the number of mice that harboured this ASV with a positive IgA probability; parentheses shows this number for taxa shaded in ‘Probability’ only. In ‘Probability’ the shading shows the 10 ASVs with the greatest sum positive IgA probability. For each ASV, their taxonomic identification is shown and for each genus thus identified other con-generic species’ ASVs are also shown. ‘Other’ shows for the other con-generic species’ ASVs the number of mice that harboured this ASV with a positive IgA probability.

| Genus | Species | ASV | Common | Probability | Other |
| --- | --- | --- | --- | --- | --- |
| *Pseudomonas* | *aeruginosa* | 33 | 24 |  |  |
|  | *fragi* | 258 |  |  | 1 |
|  | *veronii* | 353 | 10 |  |  |
|  | *aeruginosa* | 397 |  |  | 2 |
| *Alcaligenes* | *faecalis* | 52 | 22 |  |  |
|  | *aquatilis* | 446 |  |  | 0 |
| *Aquamicrobium* | NA | 57 | 22 |  |  |
| *Bacillus* | *halotolerans* | 73 | 18 |  |  |
|  | *thuringiensis* | 289 |  |  | 0 |
| *Escherichia-Shigella* | *coli* | 24 | 16 |  |  |
|  | N/A | 371 |  |  | 4 |
| *Enterococcus* | *faecium* | 71 | 16 |  |  |
|  | N/A | 345 |  |  | 2 |
| *Cutibacterium* | *acnes* | 334 | 14 |  |  |
| *Staphylococcus* | *aureus* | 163 | 11 |  |  |
|  | *equorum* | 21 |  |  | 2 |
|  | *lentus* | 81 |  |  | 3 |
|  | *saprophyticus* | 159 |  |  | 5 |
|  | *sciuri* | 257 |  |  | 1 |
| *Limosilactobacillus* | N/A | 153 | 10 |  |  |
|  | N/A | 58 |  |  | 3 |
|  | N/A | 85 |  |  | 0 |
|  | *N/A* | 92 |  |  | 3 |
| *Salmonella* | *enterica* | 135 | (9) |  |  |
| *Listeria* | *monocytogenes* | 168 | (8) |  |  |

### 3.4 The eukaryotic microbiome

We resolved 2,666 eukaryotic ASVs across both the faecal and caecal samples, which represented 32 phyla, with 6 being known gut residents; specifically, in order of relative faecal and caecal abundance combined: Ascomycota, Basidiomycota, nematode, Apicomplexa, Bigyra and Ciliophora (**Supplementary Information 15**). The mean caecal and faecal prevalence was 100% for Ascomycota and Basidiomycota, 88% for the Apicomplexa, 66% for Nematoda, 65% for Ciliophora, and 11% for Bigyra (**Supplementary Information 15**).

Mice from the three different sample sites varied in the relative abundance of these phyla in the caecal and faecal microbiome. Mice from the Nottingham site had nematode and apicomplexan infection (nematode caecal and faecal 64.9 and 1.5%, respectively; Apicomplexa caecal and faecal 9.7 and 19.2%, respectively), but these were essentially absent from Southport and Wirral mice (**Supplementary Information 16**). Consequently, the relative abundance of the caecal Ascomycota and Basidiomycota is comparatively lower in the Nottingham-derived mice, as too the faecal Basidiomycota (**Supplementary Information 16**).

Focussing on the nematode and apicomplexan infections that were concentrated in the Nottingham-derived mice, the genus identity of these ASVs showed that the nematodes were principally *Trichuris* and of the Oxyurid (pinworm) order, and that the Apicomplexa were *Eimeria* (see Section 3.6, below).

We investigated factors affecting the alpha diversity of the eukaryotic microbiome, testing for effects stepwise as: (i) mouse traits, (ii) immune-related traits, (iii) gut inflammation, (iv) the faecal eukaryotic microbiome, (v) the caecal eukaryotic microbiome and (vi) the main bacterial phyla; for beta diversity we used a PERMANOVA analysis. This showed that alpha diversity of the caecal and faecal eukaryotic microbiome was affected by: (a) one gut inflammation trait, specifically small intestine inflammation; (b) a number of caecal and faecal eukaryotic microbiome traits, where the caecal Apicomplexa and Ascomycota affected both the caecal and faecal eukaryotic alpha diversity; and (c) one bacterial microbiome trait, specifically that Actinobacteriota affected the faecal eukaryotic microbiome alpha diversity (**Supplementary Information 17**).

The beta diversity was affected by: (a) mouse traits, with sample site affecting both caecal and faecal beta diversity, and BMI affecting caecal beta diversity; (b) one immune-related trait, [faecal IgA] affecting faecal beta diversity; (c) at least 8 eukaryotic microbiome traits – Apicomplexa (faecal), Ascomycota (caecal and faecal), Basidiomycota (caecal and faecal), nematode (caecal and faecal), faecal eukaryotic microbiome alpha diversity – which all affected both caecal and faecal beta diversity, and with caecal eukaryotic microbiome alpha diversity also affecting caecal beta diversity; and (d) two bacterial microbiome traits, Proteobacteria and Actinobacteriota affecting caecal and faecal beta diversity, respectively (**Supplementary Information 18**).

It is of note that both the alpha and beta diversity of the eukaryotic microbiome is affected by bacterial taxa, which is analogous to the effects of eukaryotic taxa on the bacterial microbiome’s beta diversity (Section 3.2, above).

### 3.5 Bacterial – Eukaryotic interactions

Comparison of the factors that affect the beta diversity of the bacterial microbiome and the alpha and beta diversity of the caecal and faecal eukaryotic microbiome (**Supplementary Information 9, 18, 19**) show that three taxa have common effects: Basidiomycota, Ascomycota, and nematodes. The presence of these eukaryotic taxa therefore seem to have wide-ranging effects on the eukaryotic microbiome, but also on the bacterial microbiome. A fourth eukaryotic taxa, the Apicomplexa, also has wide-ranging effects on the eukaryotic microbiome, affecting the alpha and beta diversity of the caecal and faecal eukaryotic microbiome.

We identified 12 bacterial ASVs that more commonly bound by IgA (Table 1, section 3.3) than other bacterial ASVs. Combined, these 12 ASVs did not have a significant effect on the faecal bacterial, caecal eukaryotic, and faecal eukaryotic beta diversity (**Supplementary Information 9 and 19**).

We constructed networks of the 75 most abundant bacterial and eukaryotic taxa (**Supplementary Information 20**). These shows that most of the edges among taxa nodes are bacterial-bacterial or eukaryotic-eukaryotic, that there are some (8 positive and 9 negative) that are bacterial-eukaryotic; specifically. In the whole dataset network, *Eimeria* was within large cluster of mainly bacterial taxa, and had both positive and negative associations with bacterial taxa. In the samples site-specific networks, *Eimeria* was only present in the Nottingham network, and then with only associations to other eukaryotes.

### 3.6 *Eimeria* infection

Given the ASV identification of the Apicomplexa being principally *Eimeria*, we specifically tested for this infection by PCR. We detected *Eimeria* infection in both caecal and faecal samples of mice, with a strong positive association between diagnosis in the two (Pearson’s Chi-squared test: Χ^*2*^_1_ = 33.55, *p* <0.001), though detection was more likely in faecal samples (McNemar’s Chi-squared test: Χ^2^_1_ = 4.14, *p* =0.041); hereafter we report only faecal data. Overall, *Eimeria* was a common infection with a prevalence of 69%, though this varied among sample sites. Specifically, there was a 97% prevalence in Nottingham mice, which was significantly greater than in Southport and the Wirral (20 and 58%, respectively) (Fisher’s exact test, both *p* < 0.001 and *p* = 0.008, respectively); the non-Nottingham sites’ prevalence did not differ significantly (Fisher’s exact test, *p* = 0.057).

We investigated whether *Eimeria* infection was associated with aspects of gut immunology and physiology. We found that [faecal IgA] was significantly higher in *Eimeria*-positive than *Eimeria*-negative mice (mean ± SE: 705 ± 118 and 302 ± 106 mg/g faeces, respectively; Wilcox two-sample test: W = 170, *p* = 0.001). There was also a significantly higher [mucin] in *Eimeria*-positive mice than *Eimeria*-negative mice (mean ± SE: 9.6 ± 1.6 and 9.3 ± 2.7 mg/g faeces, respectively; GLMM Wald’s *X*2 = 6.40, *p* = 0.011). Despite these associations, there was no significant difference in inflammation in the caecum or small or large intestine between *Eimeria*-positive mice and *Eimeria*-negative mice.

The beta diversity of the bacterial microbiome, and the alpha and beta diversity of the faecal and caecal eukaryotic microbiome are affected by Apicomplexa (**Supplementary Information 9, 18 and 19**), which may be principally an effect of *Eimeria* infection. We specifically tested for this: (i) for alpha diversity by comparing models that included Apicomplexa (**Supplementary Information 18**) where it was replaced by *Eimeria*, finding that the model results were essentially unchanged; and (ii) for beta diversity with a PERMANOVA analysis finding that *Eimeria* faecal abundance affected eukaryotic (caecal and faecal) and bacterial beta diversity, and *Eimeria* caecal abundance affected eukaryotic (caecal) beta diversity (**Supplementary Information 19**).

## 4. Discussion

Our overall aim was to describe the bacterial and eukaryotic microbiome of wild mice and to determine how its composition was affected by aspects of animals’ biology. The bacterial microbiome is dominated by taxa from the Bacteroidota and Firmicutes phyla, as is almost universally observed in studies of wild and laboratory mice, and other animals. However, we find that mice vary very substantially in the relative proportion of these phyla. Analogously, for the caecal and faecal eukaryotic microbiome we found a wide range of taxa, principally from 6 phyla of know gut residents, but which was dominated by Ascomycota and Basidiomycota. Both the bacterial and eukaryotic microbiome’s beta diversity, but not alpha diversity, was affected by the site the mice came from.

We measured a range of traits, and their correlations, of the mice to seek to understand how these might affect the microbiome composition. Among these was an association of the [faecal IgA] and principal components that include age and BMI, which is consistent with age and condition-associated accumulation of antibodies in wild mice, as previously observed [15]. There were also associations of a number of aspects of the bacterial (proportion of IgA^+^) and eukaryotic (Ascomycota, Basidiomycota, nematode) microbiome with the first principal component. This is suggestive of these components of the microbiome either promoting the faecal IgA response and /or being affected by it. It is notable that there are correlations among bacterial and eukaryotic components of the microbiome, which is suggestive of potential interactions between taxa in these different groups.

We measured the abundance of eukaryotic taxa in the caecum and in faeces and find for the nematodes, Ascomycota and Basidiomycota that these are negatively correlated. This suggests that faecal measurement of these taxa (which is commonly done) is a poor measure of their abundance in the gut. Our results are also similar to studies of wild *Mus musculus* and three species of wild rodents (*Apodemus speciosus, A. argenteus, Myodes rufocanus*) which found considerable differences in the microbiome between the small intestine and the lower gut [48–51].

Considering the variation among individual mice, our PCA analysis showed that there is great variation among them though they generally group by the site from which they came. These among-site differences are consistent with other observations of differences in among-population bacterial microbiome variation [1,3,5]. Similar phenomena have been reported for *A. sylvaticus*, where mother–offspring microbiomes are idiosyncratic, before becoming more similar among mice with age [12]. Our results then provide a perspective at a larger scale, showing the microbiome diversity among mice from different geographical regions, but also among mice within sample sites.

We then investigated what affected the bacterial and eukaryotic diversity. For alpha diversity, the bacterial microbiome was only affected by itself (Bacteroidota abundance), but the eukaryotic microbiome’s was affected by microbiome composition both eukaryotic (Apicomplexa and Ascomycota) and bacterial (Actinobacteriota), as well as a single main trait of gut inflammation. In contrast, for beta diversity the bacterial microbiome was affected by mouse traits (site and age), and bacterial and eukaryotic microbiome composition; the eukaryotic microbiome by mouse traits (sample site, BMI, [faecal IgA]) and eukaryotic and bacterial microbiome composition. Moreover, the Basidiomycota, Ascomycota and nematode components of the eukaryotic microbiome affect both the bacterial and eukaryotic diversity. These observations are consistent with previous reports of the importance of fungal taxa on microbiome biology, especially in non-laboratory mice [9–10]. Nematodes are large metazoans and may therefore bring about wide-ranging effects on the microbial microbiome. Gut inflammation and immune-related traits have limited effects on the microbiome diversity, with only small intestine inflammation affecting eukaryotic microbiome alpha diversity and the [faecal IgA] affecting eukaryotic microbiome beta diversity. Instead, the principal affecters of bacterial and eukaryotic microbiome diversity are the bacterial and eukaryotic microbiomes themselves, both within and between the bacterial and eukaryotic compartments.

Therefore, while the biology of the host can affect the diversity of the microbiome we find more far-reaching effects of the microbiome composition itself, both within-bacterial and within-eukaryotic, as well as cross bacterial-eukaryotic. The evidence of an interplay between bacterial and eukaryotic taxa that we have found emphasises that studies of the bacterial microbiome should not be undertaken in isolation from a wider consideration of the gut microbiome. Throughout these analyses there are a number of indirect effects between the bacterial and eukaryotic microbiome. We assayed this directly using a network analysis of the 75 most abundant ASVs. This showed that while the network was dominated by bacterial-bacterial and eukaryotic-eukaryotic links, there were a number of bacterial-eukaryotic links.

The multiple taxa in a microbiome can induce immune responses and other host physiological changes that could affect the microbiome composition. We measured host IgA production and gut inflammation but we could not detect any effect of these on the bacterial and eukaryotic microbiome except for gut inflammation and of [faecal IgA] affecting eukaryotic alpha and beta diversity, respectively.

Among the eukaryotic microbiome was the Apicomplexan *Eimeria*, which is a well-known pathogen and it commonly infected our mice, particularly those from Nottingham where the prevalence was 97%. We found direct effects of *Eimeria* (and Apicomplexa) on eukaryotic alpha and beta diversity and on bacterial beta diversity. The network analysis showed *Eimeria* was within a cluster containing bacterial and eukaryotic taxa and that it had both positive and negative associations with some bacterial taxa. This therefore shows how a single eukaryotic taxon can have far-reaching effects on the composition of the microbiome. *Eimeria* infection affects the [faecal IgA] and of [mucin], indicative of *Eimeria*’s pathogenesis, though we could not detect direct effects on gut inflammation that might have been expected. Analysis of the gut inflammation in the three different gut sites shows no correlation, suggesting that any gut inflammation that occurs is localised. In this regard *Eimeria* might cause very localised inflammation at the site of infection, but which was not captured by our histological analyses. In contrast, the [faecal IgA] may provide a more whole-gut view of immune state.

We investigated which bacterial taxa were immune-targeted by the mice by identifying those that were IgA^+^. We find that a 12% of the microbiome is IgA^+^. In laboratory mice a range of proportions of gut bacteria being bound by IgA have been reported, from a low of 7.4 to *c*.20% [20,52]; in people a range of values for the proportion of faecal bacteria that are bound by IgA have also been reported, for example *c*.12-15, 40, 10-50, 2.5-35% [53–55]. Wild mice have higher average [faecal IgA] than do laboratory mice [14] and so it is interesting that the proportion of their bacteria bound to IgA is within the range reported for laboratory mice.

Our results show that wild mice generate IgA that binds to a minority of bacterial taxa that they harbour, and while there is some commonality in the ASVs to which IgA responses are directed these are not universal and indeed mice are also quite idiosyncratic in the ASVs they target with IgA. Similar phenomena have been reported in analysis of IgA^+^ bacteria in human faeces [53]. Mice also differ substantially in the amount of IgA binding that they undertake, with approximately a third of mice responsible for approximately half of all ASVs bound with IgA. This is consistent with the idea that mice are quite distinct in the quantity and quality of their gut IgA responses, likely reflecting their particular infection history. The mouse IgA responses are highly specific, shown by IgA targeting certain bacterial ASVs, while apparently ignoring con-generic ASVs. This among-mouse heterogeneity also means that the IgA environment experienced by an ASV, which will act as a selective pressure against the bacteria [21], is not uniform for the bacterial population distributed among different individual host mice. These findings of substantial among-individual mouse heterogeneity in the IgA binding of gut bacteria is analogous to our finding of considerable idiosyncrasy in the microbiome composition, for example the highly variable proportion of the microbiome that is Bacteroidota *vs*. Firmicutes. It is also analogous to previous reports of among-individual variation in microbiome composition [1].

## 5. Conclusion

Our work has described the composition of the bacterial eukaryotic microbiome, finding that it varies substantially among individual mice. Our analysis of the bacterial taxa that mice target by IgA shows that there is a common set of bacteria taxa targeted in this way, but also that mice are idiosyncratic in this regard. The diversity and composition of the microbiome is affected by a few specific mouse traits, but more commonly by other components of the microbiome. We detected frequent effects between components of the bacterial and eukaryotic microbiome. While we describe these effects, the mechanism by which they occur remains to be understood. The results of our work emphasise that wild animal microbiomes are distinct from those of laboratory animals, and the richness of the gut microbiome ecology deserves further investigation.

## Competing interests

The authors declare that no competing interests exist.

## Acknowledgements

This work was supported by the University of Liverpool; SHB was supported by a NERC ACCE studentship. We would like to thank Amanda Davidson for technical support; John Cameron, Stephen Hoyland, Irene Doherty for helpful access to field sites; Emanuel Heitlinger for *Eimeria* oocysts; Kathryn Else for providing faeces from IgMi mice.

## CRediT author contributions

**Louise Cheynel**: Conceptualization, Data curation, Formal analysis, Methodology, Visualization, Writing – original draft

**Simon Hunter-Barnett**: Conceptualization, Data curation, Formal analysis, Methodology, Visualization, Writing – original draft

**Lukasz Lukomski**: Methodology.

**Chris Law**: Methodology.

**Lorenzo Ressel**: Methodology.

**Jane Hurst**: Conceptualization; Supervision

**Mark Viney**: Conceptualization, Formal analysis, Funding acquisition, Project administration, Supervision, Writing – original draft the paper.

## Notes

### Competing Interest Statement

The authors have declared no competing interest.

## References

1. Hanski, E., Joseph, S., Curtis, M.A., Swann, J.W., Vallier, M., Linnerbrink, M. et al. (2025) Wild house mice have a more dynamic and aerotolerant gut microbiota than laboratory mice. BMC Microbiology, 25, 204.

2. Kreisinger, J., Cížková, D., Vohánka, J., Piálek, J. (2014) Gastrointestinal microbiota of wild and inbred individuals of two house mouse subspecies assessed using high-throughput parallel pyrosequencing. Molecular Ecology, 23, 5048–5060.

3. Suzuki, T.A., Phifer-Rixey, M., Mack, K.L., Sheehan, M.J., Lin, D., Bi, K. et al. (2019) Host genetic determinants of the gut microbiota of wild mice. Molecular Ecology, 28, 3197–3207.

4. Schmidt, E., Mykytczuk, N., Schulte-Hostedde, A.I. (2019) Effects of the captive and wild environment on diversity of the gut microbiome of deer mice (*Peromyscus maniculatus*). The ISME Journal, 13, 1293–1305.

5. Marsh, K.J., Raulo, A.M., Brouard, M., Troitsky, T., English, H.M., Allen, B. et al. (2022) Synchronous seasonality in the gut microbiota of wild mouse populations. Frontiers in Microbiology, 13, 809735.

6. Scholier, T., Lavrinienko, A., Kallio, E.R., Watts, P.C., Mappes, T. (2024) Effects of past and present habitat on the gut microbiota of a wild rodent. Proceedings of the Royal Society B, 291, 20232531.

7. Rosshart, S.P., Vassallo, B.G., Angeletti, D., Hutchinson, D.S., Morgan, A.P., Takeda, K. et al. (2017) Wild mouse gut microbiota promotes host fitness and improves disease resistance. Cell, 171, 1015–1028.e13.

8. Bruno, P., Schüler, T., Rosshart, S.P. (2025) Born to be wild: utilizing natural microbiota for reliable biomedical research. Trends in Immunology, 46, 17–28.

9. Leung, J.M., Budischak, S.A., Chung The, H., Hansen, C., Bowcutt, R., Neill, R. et al. (2018) Rapid environmental effects on gut nematode susceptibility in rewilded mice. PLoS Biology, 16, e2004108.

10. Yeung, F., Chen, Y.H., Lin, J.D., Leung, J.M., McCauley, C., Devlin. J.C. (2020) Altered immunity of laboratory mice in the natural environment is associated with fungal colonization. Cell Host and Microbe, 27, 809–822.

11. Raulo, A., Bürkner, PC., Finerty, G.E., Dale, J., Hanski, E., English, H.M. et al. (2024) Social and environmental transmission spread different sets of gut microbes in wild mice. Nature Ecology and Evolution, 8, 972–985.

12. Wanelik, K.M., Raulo, A., Troitsky, T., Husby, A. Knowles, S.C.L. (2023) Maternal transmission gives way to social transmission during gut microbiota assembly in wild mice. Animal Microbiome, 5, 29.

13. Beura, L.K., Hamilton, S.E., Bi, K., Schenkel, J.M., Odumade, O.A., Casey, K.A. et al. (2016) Normalizing the environment recapitulates adult human immune traits in laboratory mice. Nature, 532, 512–516.

14. Abolins, S., King, E.C., Lazarou, L., Weldon, L., Hughes, L., Drescher, P. et al. (2017) The comparative immunology of wild and laboratory mice, Mus musculus domesticus. Nature Communications, 8, 14811.

15. Abolins, S., Lazarou, L., Weldon, L., Hughes, L., King, E.C., Drescher, P. et al. (2018) The ecology of immune state in a wild mammal, Mus musculus domesticus. PLoS Biology, 16, e2003538.

16. Viney, M., Riley, E.M. (2017) The immunology of wild rodents: current status and future prospects. Frontiers in Immunology, 8, 1481.

17. Cheynel, L., Lazarou, L., Riley, E.M., Viney, M. (2023) The genetics of immune and infection phenotypes in wild mice, Mus musculus domesticus. Molecular Ecology, 32, 4242–4258.

18. Rosshart, S.P., Herz, J.H., Vassallo, B.G., Hunter, A., Wall, M.K., Badger, J.H. et al. (2019) Laboratory mice born to wild mice have natural microbiota and model human immune responses. Science, 365, eaaw4361.

19. Ma, J., Urgard, E., Runge, S., Classon, C.H., Mathä, L., Stark, J.M. et al. (2023) Laboratory mice with a wild microbiota generate strong allergic immune responses. Science Immunology, 8, eadf7702.

20. Palm, N. W., De Zoete, M. R., Cullen, T. W., Barry, N. A., Stefanowski, J., Hao, L. et al. (2014) Immunoglobulin A coating identifies colitogenic bacteria in inflammatory bowel disease. Cell, 158, 1000–1010.

21. Viney, M., Cheynel, L. (2023) Gut immune responses and evolution of the gut microbiome – a hypothesis. Discovery Immunology, 2, kyad025.

22. Laforest-Lapointe, I., Arrieta, M.C. (2018). Microbial eukaryotes: a missing link in gut microbiome studies. MSystems, 3, e00201–17.

23. Liao, Y., Gao, I.H., Kusakabe, T., Lin, W-Y., Grier, A., Pan, X. et al. (2024) Fungal symbiont transmitted by free-living mice promotes type 2 immunity. Nature, 636, 697–704.

24. Labocha, M.K., Schutz, H., Hayes, J.P. (2014) Which body condition index is best? Oikos, 123, 111–119.

25. Rowe, F.P., Bradfield, A., Quy, R.J., Swinney, T. (1985) Relationship between eye lens weight and age in the wild house mouse (Mus musculus). Journal of Applied Ecology, 22, 55–61.

26. Williams, J.M., Duckworth, C.A., Vowell, K., Burkitt, M.D., Pritchard, D.M. (2016) Intestinal preparation techniques for histological analysis in the mouse. Current Protocols in Mouse Biology, 6, 148–168.

27. McGuckin, M.A., Lindén, S.K., Sutton, P., Florin, T.H. (2011) Mucin dynamics and enteric pathogens. Nature Reviews Microbiology, 9, 265–278.

28. Ansia, I., Drackley, J.K. (2020). Technical note: Evaluation of 3 methods to determine mucin protein concentration in ileal digesta of young preweaning calves. Journal of Dairy Science, 103, 6250–6257.

29. Crowther, R.S., Wetmore, R.F. (1987) Fluorometric assay of O-linked glycoproteins by reaction with 2-cyanoacetamide. Analytical Biochemistry, 163, 170–174.

30. Hunter-Barnett, S. (2023) The gut eukaryome of wild house mice. Doctor of Philosophy thesis, University of Liverpool.

31. Erben, U., Loddenkemper, C., Doerfel, K., Spieckermann, S., Haller, D., Heimesaat, M.M. et al. (2014) A guide to histomorphological evaluation of intestinal inflammation in mouse models. International Journal of Clinical and Experimental Pathology, 7, 4557–4576.

32. Duszynski, D.W. (2021) Biodiversity of the Coccidia (Apicomplexa: Conoidasida) in vertebrates: what we know, what we do not know, and what needs to be done. Folia Parasitolgica, 68, 001.

33. Hunter-Barnett, S., Viney, M. (2024) Gut protozoa of wild rodents - a meta-analysis. Parasitology 1–12, 10.1017/S0031182024000556.

34. Vandeputte, D., Kathagen, G., D’hoe, K., Vieira-Silva, S., Valles-Colomer, M., Sabino, J. et al. (2017) Quantitative microbiome profiling links gut community variation to microbial load. Nature, 551, 507–511.

35. Caporaso, J.G., Lauber, C.L., Walters, W.A., Berg-Lyons, D., Huntley, J., Fierer, N. et al. (2012) Ultra-high-throughput microbial community analysis on the Illumina HiSeq and MiSeq platforms. ISME Journal, 6, 1621–4.

36. Kim, D., Hofstaedter, C.E., Zhao, C., Mattei, L., Tanes, C., Clarke, E. et al. (2017). Optimizing methods and dodging pitfalls in microbiome research. Microbiome, 5, 52.

37. Mann, A.E., Mazel, F., Lemay, M.A., Morien, E., Billy, V. et al. (2020) Biodiversity of protists and nematodes in the wild nonhuman primate gut. The ISME Journal, 14, 609–622.

38. Dray, S., Dufour, A.B. (2007) The ade4 package: implementing the duality diagram for ecologists. Journal of Statistical Software, 22, 1–20.

39. Jackson, M.A., Pearson, C., Ilott, N.E., Huus, K.E., Hegazy, A.N., Webber, J. et al. (2021) Accurate identification and quantification of commensal microbiota bound by host immunoglobulins. Microbiome, 9, 1–22.

40. Lahti, L., Shetty, S. (2017) Tools for microbiome analysis in R. R package version 1.19.1. https://microbiome.github.io/tutorials/

41. Burnham, K.P., Anderson, D.R. (2002) Model selection and multimodel inference: a practical information-theoretic approach. New York, NY: Springer New York.

42. McMurdie, P.J., Holmes, S. (2013) phyloseq: An R Package for reproducible interactive analysis and graphics of microbiome census data. PLoS One, 8, e61217.

43. Cao, Q., Sun, X., Rajesh, K., Chalasani, N., Gelow, K., Katz, B. et al. (2021) Effects of rare microbiome taxa filtering on statistical analysis. Frontiers in Microbiology, 11, 607325.

44. Peschel, S., Müller, C.L. von Mutius, E., Boulesteix, A-L., Depner, M. (2020) NetCoMi: network construction and comparison for microbiome data in R. Briefings in Bioinformatics, 22, bbaa290.

45. Yoon G., Gaynanova I., Müller, C.L. (2019) Microbial networks in SPRING - semi-parametric rank-based correlation and partial correlation estimation for quantitative microbiome data. Frontiers in Genetics, 10, 516.

46. Holm S. (1979). A simple sequentially rejective multiple test procedure. Scandinavian journal of statistics, 65–70.

47. Bates, D., Maechler, M., Bolker, B., Walker, S., Christensen, R.H.B., Singmann, H. et al. (2015). Package ‘lme4’. Convergence, 12, 2.

48. Gu, S., Chen, D., Zhang, J.N., Lv, X., Wang, K., Duan, L.P. et al. (2013) Bacterial community mapping of the mouse gastrointestinal tract. PLoS One, 8, e74957.

49. Suzuki, T.A., Nachman, M.W. (2016) Spatial heterogeneity of gut microbial composition along the gastrointestinal tract in natural populations of house mice. PLoS One, 11, e0163720.

50. Anders, J.L., Moustafa, M.A.M., Mohamed, W.M.A., Hayakawa, T., Nakao, R., Koizumi, I. (2021) Comparing the gut microbiome along the gastrointestinal tract of three sympatric species of wild rodents. Scientific Reports, 11, 19929.

51. Bendová, B., Mikula, O., Vošlajerová Bímová, B., Čížková, D., Daniszová, K., Ďureje, L. et al. (2022) Divergent gut microbiota in two closely related house mouse subspecies under common garden conditions. FEMS Microbiology Ecology, 98, 1–13.

52. Bunker, J.J., Flynn, T.M., Koval, J.C., Shaw, D.G., Meisel, M., McDonald, B.D. et al. (2015) Innate and adaptive humoral responses coat distinct commensal bacteria with immunoglobulin A. Immunity, 43, 541–53.

53. Olm, M.R., Spencer, S.P., Takeuchi, T., Silva, E.L., Sonnenburg, J.L. (2025) Metagenomic immunoglobulin sequencing reveals IgA coating of microbial strains in the healthy human gut. Nature Microbiology, 10, 112–125.

54. Augustine, T., Murugesan, S., Badri, F., Gentilcore, G., Grivel, J-C., Akobeng, A. et al. (2024) Immunoglobulin-coating patterns reveal altered humoral responses to gut bacteria in pediatric cow milk allergies. Journal of Translational Medicine, 22, 1021.

55. Van Gogh, M., Louwers, J.M., Celli, A., Gräve, S., Viveen, M.C., Bosch, S. et al. (2024) Next-generation IgA-SEQ allows for high-throughput, anaerobic, and metagenomic assessment of IgA-coated bacteria. Microbiome, 12, 211.

