## Supplementary Information for "Gut immunity and the bacterial and eukaryotic microbiome of wild house mice"

**Supplementary Information 1.** The distribution of individuals by their ID, site at which they were caught, sex, age in weeks and BMI. The three sites are Nottinghamshire (52° 54' 38" N, 1° 05' 21" W); Southport, Merseyside (53° 40' 22" N, 2° 54' 24" W); the Wirral Peninsula (53° 17' 27" N, 3° 02' 00" W). In calculating body mass, for female mice that were later found to be pregnant the mass of the foetuses were subtracted from their final mass, and these values were used in all subsequent analyses.

| ID | Site | Sex | Age | BMI |
|---|---|---|---|---|
| N01 | Nottingham | F | 12.16 | 1.154 |
| N02 | Nottingham | M | 15.08 | 1.354 |
| N03 | Nottingham | M | 12.22 | 1.335 |
| N04 | Nottingham | F | 12.73 | 1.487 |
| N05 | Nottingham | M | 5.32 | 1.194 |
| N06 | Nottingham | F | 5.92 | 1.291 |
| N07 | Nottingham | M | 8.1 | 1.318 |
| N08 | Nottingham | M | 5.22 | 1.279 |
| N09 | Nottingham | M | 7.56 | 1.238 |
| N10 | Nottingham | M | 2.35 | 1.120 |
| N11 | Nottingham | F | 4.27 | 1.260 |
| N12 | Nottingham | M | 4.87 | 1.265 |
| N13 | Nottingham | F | 4.56 | 1.254 |
| N14 | Nottingham | M | 1.7 | 1.041 |
| N15 | Nottingham | F | 7.18 | 1.268 |
| N16 | Nottingham | M | 1.81 | 1.051 |
| N17 | Nottingham | M | 2.76 | 1.104 |
| N18 | Nottingham | M | 2.54 | 1.075 |
| N19 | Nottingham | M | 2.06 | 1.156 |
| N20 | Nottingham | F | 1.83 | 1.103 |
| N21 | Nottingham | M | 2.56 | 1.195 |
| N22 | Nottingham | M | 5.63 | 1.287 |
| N23 | Nottingham | M | 7.14 | 1.128 |
| N24 | Nottingham | F | 5.25 | 1.101 |
| N25 | Nottingham | M | 7.51 | 1.336 |
| N26 | Nottingham | F | 9.21 | 1.219 |
| N27 | Nottingham | F | 9.52 | 1.344 |
| N28 | Nottingham | M | 16.27 | 1.291 |
| N29 | Nottingham | F | 4.26 | 1.194 |
| N30 | Nottingham | M | 9.42 | 1.220 |
| N31 | Nottingham | M | 7.3 | 1.293 |
| S01 | Southport | M | 4.41 | 1.218 |
| S02 | Southport | F | 4.41 | 1.166 |
| S03 | Southport | M | 6.4 | 1.208 |
| S04 | Southport | M | 6.22 | 1.236 |
| S05 | Southport | M | 9.47 | 1.371 |
| S06 | Southport | M | 6.26 | 1.226 |

| | | | | |
|---|---|---|---|---|
| S07 | Southport | M | 7.22 | 1.304 |
| S08 | Southport | M | 12.16 | 1.154 |
| S09 | Southport | M | 15.08 | 1.354 |
| S10 | Southport | F | 12.22 | 1.335 |
| S11 | Southport | F | 12.73 | 1.487 |
| S12 | Southport | F | 5.32 | 1.194 |
| S13 | Southport | F | 5.92 | 1.291 |
| S14 | Southport | M | 8.1 | 1.318 |
| S15 | Southport | F | 5.22 | 1.279 |
| W01 | Wirral | M | 7.56 | 1.238 |
| W02 | Wirral | M | 2.35 | 1.120 |
| W03 | Wirral | F | 4.27 | 1.260 |
| W04 | Wirral | M | 4.87 | 1.265 |
| W05 | Wirral | M | 4.56 | 1.254 |
| W06 | Wirral | M | 1.7 | 1.041 |
| W07 | Wirral | F | 7.18 | 1.268 |
| W08 | Wirral | M | 1.81 | 1.051 |
| W09 | Wirral | M | 2.76 | 1.104 |
| W10 | Wirral | M | 2.54 | 1.075 |
| W11 | Wirral | M | 2.06 | 1.156 |
| W12 | Wirral | M | 1.83 | 1.103 |

**Supplementary Information 2**

**Supplementary Methods**

**1. Intestinal antibody response**

Faeces were dissolved in 50 μL PBS supplemented with a protease inhibitor cocktail to a final concentration of 20% v/v (SIGMAFAST Protease Inhibitor cocktail tablets, EDTA-free) to form a slurry that was left at room temperature for 1 hour. The samples were then centrifuged (13,000 x g, 10 minutes, 4°C), the supernatant removed and IgA concentration measured using a commercially available mouse IgA ELISA kit following the manufacturer's instructions (Invitrogen). Standard curves were constructed using the mouse IgA supplied with the kit, and this was used to interpolate the samples' IgA concentration as mg IgA per g of faeces.

**2. *Eimeria* infection**

Faecal and caecal DNA was extracted using the QIAamp PowerFeacl Pro DNA Kit (Qiagen). For caecal contents the protocol was adjusted to account for the samples being stored in PBS; specifically, prior to use in the kit caecal samples were centrifuged (20,000 x g, 10 minutes, 4°C) and the supernatant discarded. The contents of the PowerBead Pro Tubes and 800 μL of the kit's solution CD1 were then added to the pellet of caecal contents after which the manufacturer's instructions were followed. The DNA samples were then used in a PCR to identify the presence of *Eimeria* using the primers Ap5_Fwd (YAAAGGAATTTGAATCCTCGTTT) and Ap5_Rev (YAGAATTGATGCCTGAGYGGTC), that target a region of the apicoplast genome (Jarquín-Díaz *et al*., 2019). Positive controls were DNA from oocysts of *E. falciformis* and *E. ferrisi.* Successful amplification of *Eimeria* was defined as the presence of a ~448 bp amplicon; we triplicated the amplification of samples from each mouse and only recorded a mouse as *Eimeria* positive if all three replicates were positive.

**3. Flow cytometry counting and sorting of $IgA^+$ and $IgA^-$ faecal bacteria**

Mouse faecal pellets were incubated in 1 mL PBS per 100 mg faeces on ice for 1 hour. Faecal pellets were mechanically homogenized with a stirrer and then centrifuged (50 x g, 15 minutes, 4°C) to remove large particles, and faecal bacteria in the supernatant were removed (100 μL per sample), washed with 1 mL staining buffer (PBS with 1% w/v Bovine Serum Albumin (BSA)) and centrifuged (8,000 x g, 5 minutes, 4°C) before resuspension in 1 mL staining buffer. A 20 μL aliquot sample of this bacterial suspension was saved as the pre-sort sample for 16S sequencing analysis. The remaining bacterial suspension was first stained with 1 μL SYBR Green I (1:100 dilution in dimethylsulfoxide Thermo Fisher Scientific), for 20 minutes, in the dark, at 37 °C. After an additional wash in staining buffer bacterial pellets were resuspended in 100 μL blocking buffer (staining buffer containing 20% v/v normal rat serum) and incubated on ice for 20 minutes. Finally, the samples were stained with 100 μL staining buffer containing PE-conjugated anti-mouse IgA (1:12.5; eBioscience clone mA-6E1) for 30 minutes on ice. Samples were then washed 3 times with 1 mL staining buffer before flow cytometry sorting. We used two controls, (i) mouse faecal samples without using SYBR Green or PE-conjugated anti-mouse IgA, and (ii) faeces from IgMi mice, which do not produce or secrete IgA, but which were stained using the protocol above.

Samples were analysed and separated using a FACS Aria IIIu Cell Sorter (BD Bioscience, Oxford, UK). The gating strategy was derived from dot plots with the following hierarchy: scatter gates (SSC/FCS) under high PMTV (SSC = 900, FSC = 800) where larger remaining mammalian cells were expected to be excluded from the range, then SYBR[+] gate to exclude non-nucleated particles, and finally IgA[-] or IgA[hi] to detect and sort $IgA^-$ and $IgA^+$ bacteria, respectively (**Supplementary Information 21**). IgA[hi] and IgA[-] were defined from dot plots of pair-matched negative controls (no

IgA staining, as control (i), above) from each individual, where maximum tolerance to false positive was <0.1%. On-site data visualization, gating, and sort command were performed using FACS Diva software, version 8.1. The $IgA^+$ and $IgA^-$ bacteria were then pelleted (10,000 x g, 5 minutes, 4°C) and frozen at −80°C for future use.

To prepare DNA from the samples they were suspended in 400 µL staining buffer to which was added 250 µL 0.1 mm zirconia/silica beads (Biospec), 300 µL Lysis buffer (200 mM NaCl, 200 mM Tris, 20 mM EDTA, pH 8), 200 µL 20% w/v SDS and 500 µL 25 phenol : 24 chloroform : 1 isoamylalcohol). Samples were chilled on ice for 4 minutes and homogenized by bead beating (2 minutes bead beating, 2 minutes on ice, 2 minutes bead beating), after which the samples were centrifuged (6000 x g, 5 minutes, 4°C), and the aqueous phase transferred to a Phase Lock Gel tube (Light; 5 PRIME), to which an equal volume of phenol:chloroform:isoamylalcohol was added, and samples mixed by inversion and then centrifuged (16,100 x g, 3 minutes, room temperature). The DNA was then precipitated, the pellet air dried, and resuspended in 50 µL TE buffer.

**4. 16S rRNA sequencing and bioinformatic analyses**

Sequences were processed using the DADA2 pipeline tutorial (1.12) in R to call amplicon sequence variants (ASVs) (Callahan *et al.,* 2016). Sequences were examined for quality to determine trimming parameters, and sequences were trimmed at 210 bp (Forward) and (Reverse) and standard filtering parameters were used (maxN=0, truncQ=2, rm.phix=TRUE and maxEE=2). The average proportion of reads retained per sample at the end of the bioinformatics processing was 0.99. We assigned taxonomy to the ASVs using the Silva reference database (Silva version 138.1). Sample metadata, taxonomy tables, and an ASV abundance matrix were integrated into a phyloseq object for downstream analysis. Data contained 4,815 ASVs and 19,378,921 total reads (average 111,373.11; range 7,107 – 1,363,467). Prior to analysis an abundance filter was applied, eliminating ASVs with a total abundance of fewer than 100 reads, resulting in 2,383 ASVs. We successfully recovered the species present in our mock communities.

**5. 18S rRNA sequencing and bioinformatic analyses**

To identify taxonomically the sequence reads we used QIIME 2 2021.2 (Bolyen *et al.*, 2019). Sequence reads were trimmed using cutadapt (Martin, 2011) and paired-end reads merged, quality filtered and denoised using DADA2 to produce ASVs (Callahan *et al.*, 2016). ASVs were then aligned using mafft (Katoh *et al.*, 2002) and a phylogeny based on ASV sequence similarity created using fasttree (Price *et al.*, 2010). Sequences matching the 528F and 707R primers and their taxonomic classification were extracted from the SILVA 138 database (Quast *et al.*, 2013) to train a naïve Bayes classifier using the feature-classifier tool (Bokulich *et al.*, 2018). Taxonomy was then assigned to each ASV using the trained naïve Bayes classifier. The sequence data and host metadata were then transferred into R Studio using the qiime2R package (v0.99.6, Bisanz, 2018) for subsequent analysis. ASVs that failed to match the SILVA database at the phylum level were BLAST searched against the NCBI database (Altschul *et al.*, 1990; Sayers *et al.*, 2021) and taxonomy assigned manually using the R Phyloseq package (v1.38.0, McMurdie and Holmes, 2013).

**References**

Altschul, S.F., Gish, W., Miller, W., Myers, E.W., Lipman, D.J. (1990) Basic local alignment search tool. *Journal of Molecular Biology*, 215, 403-410.

Bisanz, J. E. (2018) Importing QIIME2 artifacts and associated data into R sessions. R package version 0.99.6. https://github.com/jbisanz/qiime2R.

Bolyen, E., Rideout, J.R., Dillon, M.R., Bokulich, N.A., Abnet, C.C., *et al*. (2019) Reproducible, interactive, scalable and extensible microbiome data science using QIIME 2. *Nature Biotechnology*, 37,852-857.

Bokulich, N.A., Kaehler, B.D., Rideout, J.R., Dillon, M., Bolyen, E., *et al*. (2018) Optimizing taxonomic classification of marker-gene amplicon sequences with QIIME 2's q2-feature-classifier plugin. *Microbiome*, 6, 90.

Callahan, B.J., McMurdie, P.J., Rosen, M.J., Han, A.W., Johnson, A.J. A., Holmes, S.P. (2016) DADA2: High-resolution sample inference from Illumina amplicon data. *Nature Methods*, 13, 581-583.

Jarquín-Díaz, V.H., Balard, A., Jost, J., Kraft, J., Dikmen, M.N., Kvičerová, J., *et al*. (2019) Detection and quantification of house mouse *Eimeria* at the species level – Challenges and solutions for the assessment of coccidia in wildlife. *International Journal for Parasitology: Parasites and Wildlife*, 10, 29-40.

Katoh, K., Misawa, K., Kuma, K., Miyata, T. (2002) MAFFT: a novel method for rapid multiple sequence alignment based on fast Fourier transform. *Nucleic Acids Research*, 30, 3059-3066.

Martin, M. (2011) Cutadapt removes adapter sequences from high-throughput sequencing reads. *EMBnet.journal*, 17, 10-12.

McMurdie, P.J., Holmes, S. (2013) phyloseq: An R Package for reproducible interactive analysis and graphics of microbiome census data. *PLoS One*, 8, e61217.

Price, M.N., Dehal, P.S., Arkin, A.P. (2010) FastTree 2 – Approximately Maximum-Likelihood Trees for Large Alignments. *PLoS One*, 5, 9490.

Quast, C., Pruesse, E., Yilmaz, P., Gerken, J., Schweer, T., Yarza, P., *et al*. (2013) The SILVA ribosomal RNA gene database project: improved data processing and web-based tools. *Nucleic Acids Research*, 41, D590-D596.

Sayers, E.W., Bolton, E.E., Brister, J.R., Canese, K., Chan, J., *et al*. (2021) Database resources of the National Center for Biotechnology Information. *Nucleic Acids Research*, 50, D20-D26.

**Supplementary Information 3.** PCA analysis showing (A) Percentage of variance explained by each axes. (B) Correlation between measured traits and the first 6 components of the PCA; C is caecal, F is faecal. Positive and negative correlations ≥ 0.50 are coloured in orange and blue, respectively. (C) Correlation matrix of measured traits. (D) Biplot of PCA analysis of the measured traits showing young (< 6 weeks) and adult (≥ 6 weeks) mice. (E) Biplot of PCA analysis of the measured traits showing male and female mice. In D and E Arrows indicate the traits' contribution to each of the first two PCs, with longer arrows denoting stronger correlations. Small points are individual mice, and large points are average values for mice at each site.

**(A)**

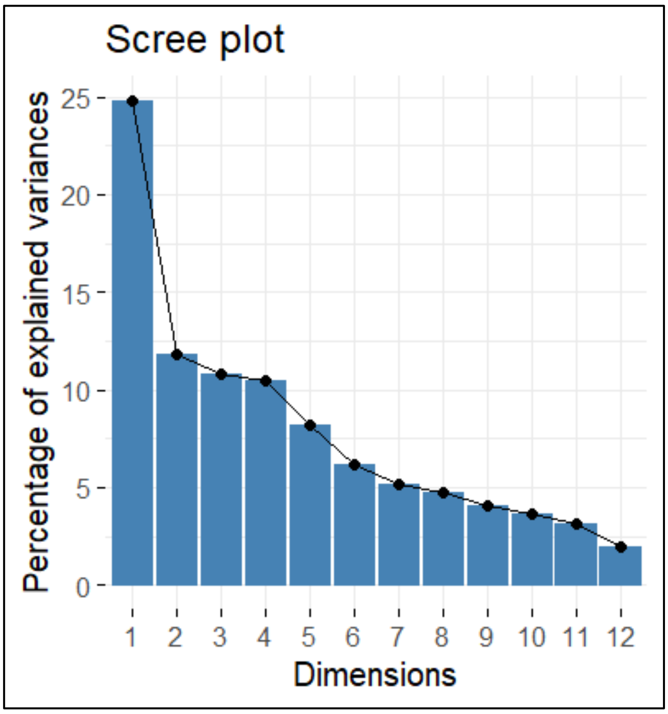

**(B)**

| | Principal Components | | | | | |
|---|---|---|---|---|---|---|
| | 1 | 2 | 3 | 4 | 5 | 6 |
| BMI | -0.53 | -0.32 | 0.58 | 0.00 | -0.02 | -0.14 |
| Age (weeks) | -0.50 | -0.35 | 0.63 | 0.00 | 0.22 | 0.07 |
| [Faecal IgA] | -0.45 | -0.62 | -0.21 | -0.05 | 0.13 | 0.15 |
| % $IgA^+$ bacteria | -0.52 | -0.41 | 0.02 | -0.01 | -0.06 | -0.25 |
| Bacterial load | 0.12 | -0.59 | -0.25 | -0.28 | -0.35 | 0.03 |
| Caecum inflammation | -0.12 | -0.48 | -0.12 | 0.37 | 0.24 | -0.39 |
| LI inflammation | -0.44 | 0.00 | -0.30 | -0.44 | 0.15 | -0.33 |
| SI inflammation | -0.01 | 0.38 | 0.03 | 0.40 | 0.52 | 0.08 |
| [Mucin] | -0.08 | 0.19 | -0.28 | -0.28 | 0.53 | -0.23 |
| Apicomplexa (C) | -0.34 | -0.26 | -0.13 | -0.17 | 0.37 | 0.69 |
| Apicomplexa (F) | -0.44 | -0.08 | -0.73 | 0.27 | -0.01 | 0.08 |
| Ascomycota (C) | 0.83 | -0.25 | 0.13 | 0.30 | 0.01 | 0.06 |
| Ascomycota (F) | -0.43 | 0.32 | 0.42 | -0.55 | -0.06 | 0.09 |
| Basidiomycota (C) | -0.79 | 0.30 | 0.03 | 0.35 | -0.16 | -0.25 |
| Basidiomycota (F) | 0.79 | -0.33 | 0.19 | 0.21 | 0.01 | -0.08 |
| Nematoda (C) | 0.61 | -0.06 | -0.13 | -0.66 | -0.04 | -0.17 |
| Nematoda (F) | -0.45 | 0.14 | -0.13 | 0.18 | -0.63 | 0.17 |

## (C)

| | BMI | Age (weeks) | [Faecal IgA] | % IgA+ bact. | Bacterial Load | Caecum Infl. | LI Infl. | SI Infl. | [Mucin] | Apicomplexa (F) | Ascomycota (F) | Basidiomycota (F) | Nematoda (F) | Apicomplexa (C) | Ascomycota (C) | Basidiomycota (C) | Nematoda (C) |
|---|---|---|---|---|---|---|---|---|---|---|---|---|---|---|---|---|---|
| BMI | | 0.70 | 0.26 | 0.32 | -0.03 | 0.15 | 0.14 | -0.11 | -0.08 | -0.10 | 0.27 | -0.19 | 0.19 | 0.09 | -0.30 | -0.26 | 0.31 |
| Age (weeks) | 0.70 | | 0.39 | 0.28 | -0.12 | 0.16 | 0.04 | -0.03 | -0.07 | -0.16 | 0.29 | -0.19 | 0.02 | 0.23 | -0.32 | -0.24 | 0.23 |
| [Faecal IgA] | 0.26 | 0.39 | | 0.40 | 0.25 | 0.22 | 0.17 | -0.14 | 0.10 | 0.40 | -0.04 | -0.22 | 0.13 | 0.33 | -0.15 | -0.21 | 0.10 |
| % IgA+ bact. | 0.32 | 0.28 | 0.40 | | 0.20 | 0.13 | 0.24 | -0.13 | 0.00 | 0.15 | 0.11 | -0.18 | 0.08 | 0.13 | -0.31 | -0.32 | 0.37 |
| Bacterial Load | -0.03 | -0.12 | 0.25 | 0.20 | | 0.13 | 0.02 | -0.27 | -0.15 | 0.03 | -0.10 | 0.17 | 0.03 | 0.09 | 0.29 | 0.04 | -0.25 |
| Caecum Infl. | 0.15 | 0.16 | 0.22 | 0.13 | 0.13 | | 0.07 | 0.01 | -0.06 | 0.24 | -0.23 | 0.05 | -0.15 | 0.02 | -0.18 | 0.06 | 0.11 |
| LI Infl. | 0.14 | 0.04 | 0.17 | 0.24 | 0.02 | 0.07 | | -0.10 | 0.2 | 0.22 | 0.17 | -0.36 | 0.05 | 0.2 | 0.07 | -0.52 | 0.21 |
| SI Infl. | -0.11 | -0.03 | -0.14 | -0.13 | -0.27 | 0.01 | -0.10 | | 0.09 | 0.01 | -0.05 | 0.00 | -0.15 | 0.02 | -0.24 | -0.03 | 0.16 |
| [Mucin] | -0.08 | -0.07 | 0.10 | 0.00 | -0.15 | -0.06 | 0.20 | 0.09 | | 0.07 | 0.00 | -0.17 | -0.12 | 0.06 | 0.13 | -0.17 | 0.00 |
| Apicomplexa (F) | -0.10 | -0.16 | 0.40 | 0.15 | 0.03 | 0.24 | 0.22 | 0.01 | 0.07 | | -0.26 | -0.47 | 0.30 | 0.21 | -0.28 | -0.32 | 0.32 |
| Ascomycota (F) | 0.27 | 0.29 | -0.04 | 0.11 | -0.10 | -0.23 | 0.17 | -0.05 | 0.00 | -0.26 | | -0.54 | 0.07 | 0.12 | 0.02 | -0.52 | 0.25 |
| Basidiomycota (F) | -0.19 | -0.19 | -0.22 | -0.18 | 0.17 | 0.05 | -0.36 | 0.00 | -0.17 | -0.47 | -0.54 | | -0.34 | -0.23 | 0.31 | 0.79 | -0.57 |
| Nematoda (F) | 0.19 | 0.02 | 0.13 | 0.08 | 0.03 | -0.15 | 0.05 | -0.15 | -0.12 | 0.30 | 0.07 | -0.34 | | -0.01 | -0.34 | -0.32 | 0.46 |
| Apicomplexa (C) | 0.09 | 0.23 | 0.33 | 0.13 | 0.09 | 0.02 | 0.20 | 0.02 | 0.06 | 0.21 | 0.12 | -0.23 | -0.01 | | -0.27 | -0.26 | -0.09 |
| Ascomycota (C) | -0.30 | -0.32 | -0.15 | -0.31 | 0.29 | -0.18 | 0.07 | -0.24 | 0.13 | -0.28 | 0.02 | 0.31 | -0.34 | -0.27 | | 0.25 | -0.73 |
| Basidiomycota (C) | -0.26 | -0.24 | -0.21 | -0.32 | 0.04 | 0.06 | -0.52 | -0.03 | -0.17 | -0.32 | -0.52 | 0.79 | -0.32 | -0.26 | 0.25 | | -0.68 |
| Nematoda (C) | 0.31 | 0.23 | 0.10 | 0.37 | -0.25 | 0.11 | 0.21 | 0.16 | 0.00 | 0.32 | 0.25 | -0.57 | 0.46 | -0.09 | -0.73 | -0.68 | |

(D)

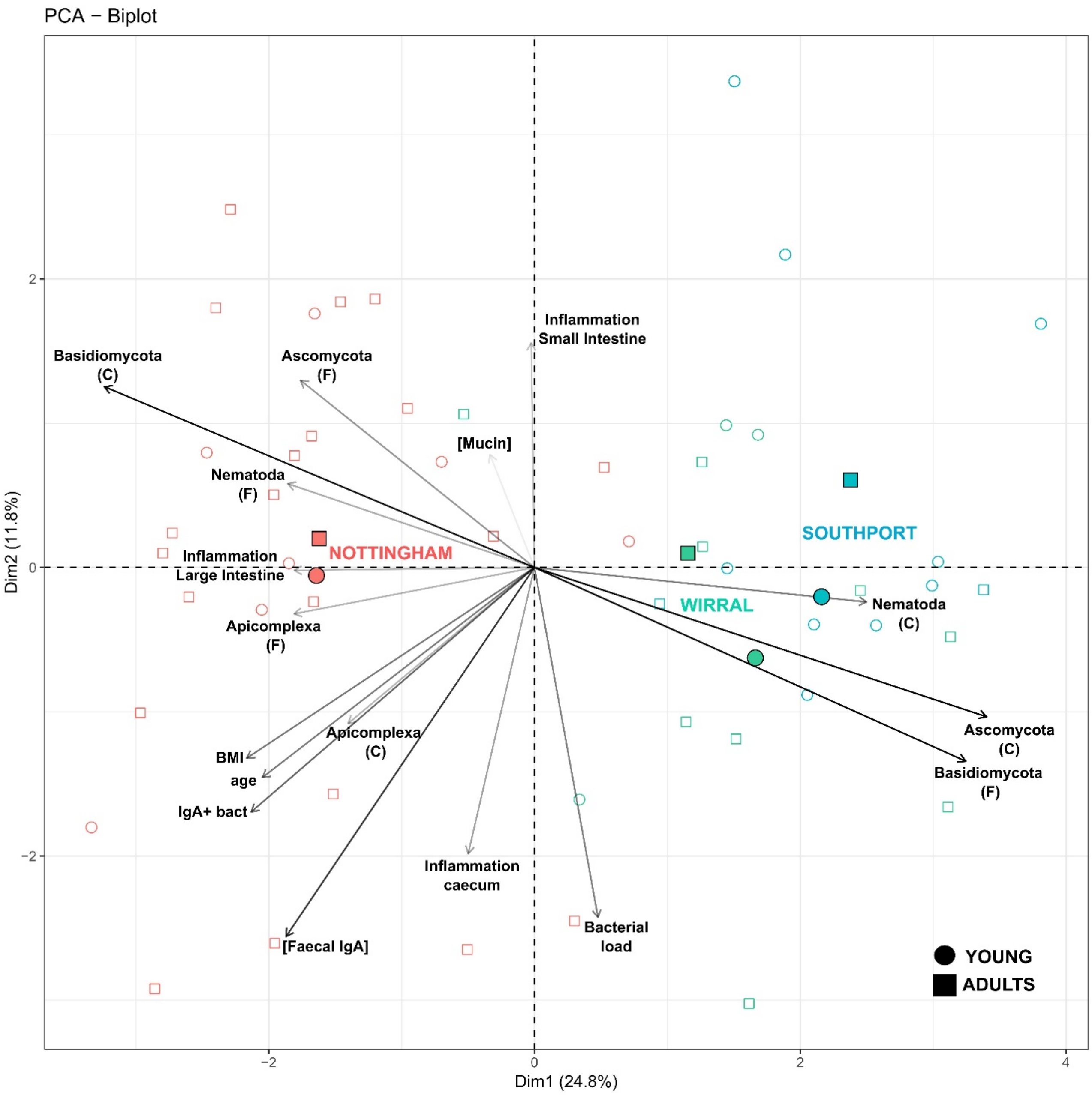

PCA − Biplot
Dim2 (11.8%)
Dim1 (24.8%)
2
0
−2
−2
0
2
4
Basidiomycota (C)
Ascomycota (F)
Inflammation Small Intestine
[Mucin]
Nematoda (F)
Inflammation Large Intestine
NOTTINGHAM
Apicomplexa (F)
Apicomplexa (C)
BMI
age
IgA+ bact
[Faecal IgA]
Inflammation caecum
Bacterial load
SOUTHPORT
WIRRAL
Nematoda (C)
Ascomycota (C)
Basidiomycota (F)
YOUNG
ADULTS

(E)

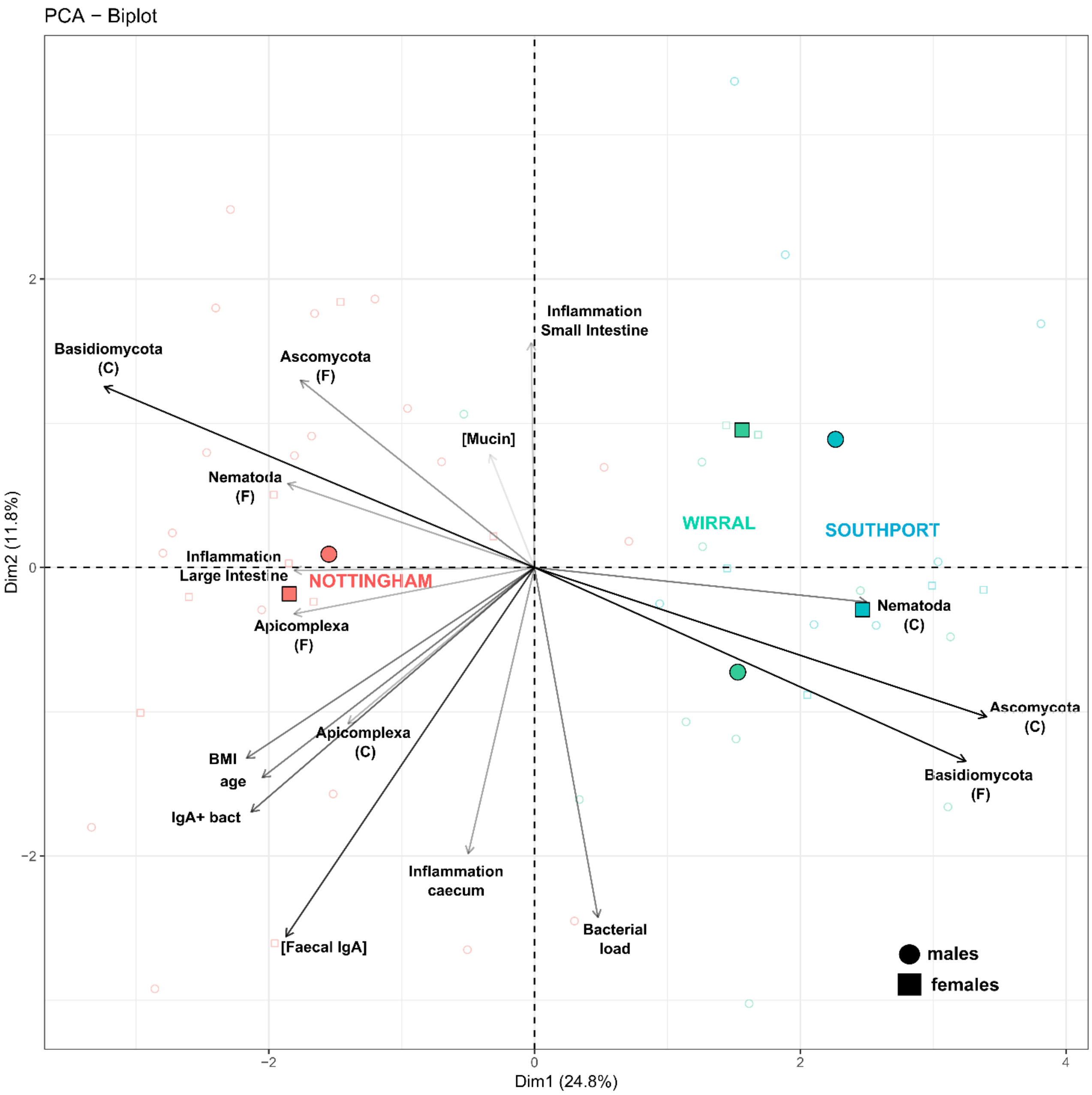
PCA – Biplot
Basidiomycota (C)
Ascomycota (F)
Inflammation Small Intestine
[Mucin]
Nematoda (F)
Inflammation Large Intestine
NOTTINGHAM
Apicomplexa (F)
WIRRAL
SOUTHPORT
Nematoda (C)
Ascomycota (C)
Basidiomycota (F)
Apicomplexa (C)
BMI
age
IgA+ bact
Inflammation caecum
Bacterial load
[Faecal IgA]
males
females
Dim1 (24.8%)
Dim2 (11.8%)

**Supplementary Information 4.** (A) [Faecal IgA] (µg / mL) as a function of mouse age in weeks. Both axes are $log_{10}$ scales. The line represents the linear relationship between [faecal IgA] and age; linear model = positive relationship and significant effect of age on [faecal IgA], $p = 0.02$. (B) [Faecal IgA] as a function of intestinal bacterial load (FACS count of number bacteria per mg of faeces). The y-axis is a $log_{10}$ scale. The line represents the linear relationship between [faecal IgA] and intestinal bacterial load; linear model = positive relationship and significant effect of bacterial load on [faecal IgA], $p < 0.001$.

**(A)**

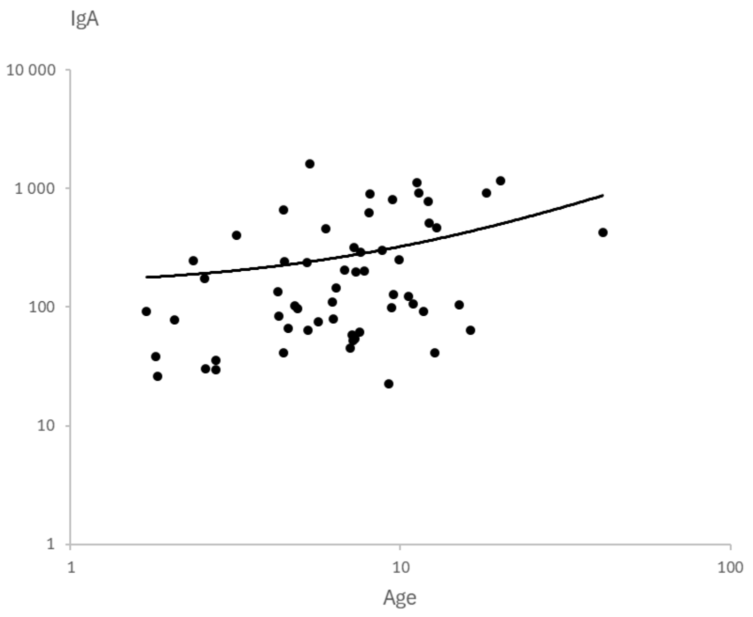

**(B)**

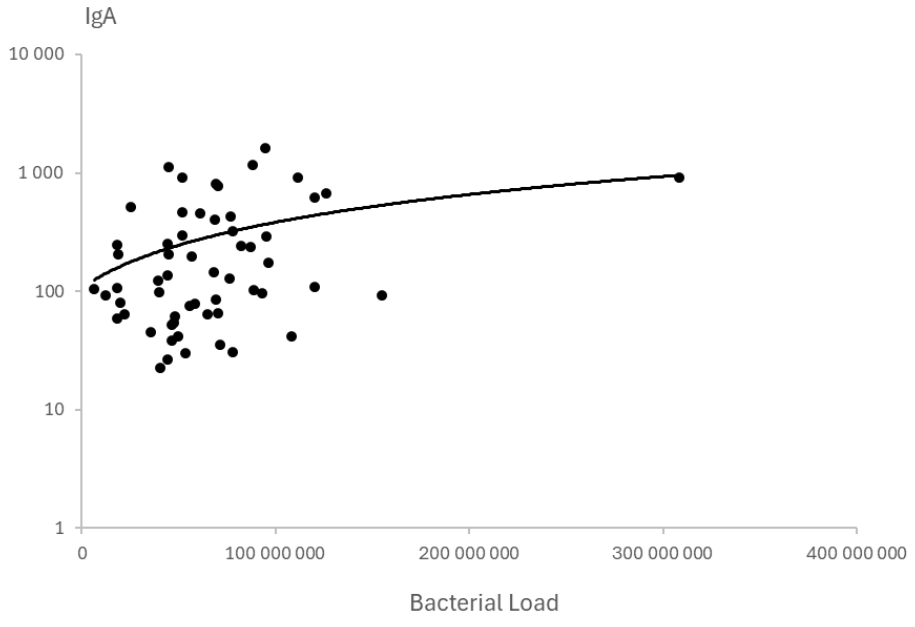

**Supplementary Information 5.** Bacterial microbiome alpha diversity calculated using Shannon's index from ASV abundance of the pre-sort samples for the three sample sites, which were compared using Wilcoxon signed-rank tests, finding that they do not differ, Kruskal-Wallis: $\chi^2_2 = 3.01$, $p > 0.05$. Whiskers show the minimum to maximum values in the dataset.

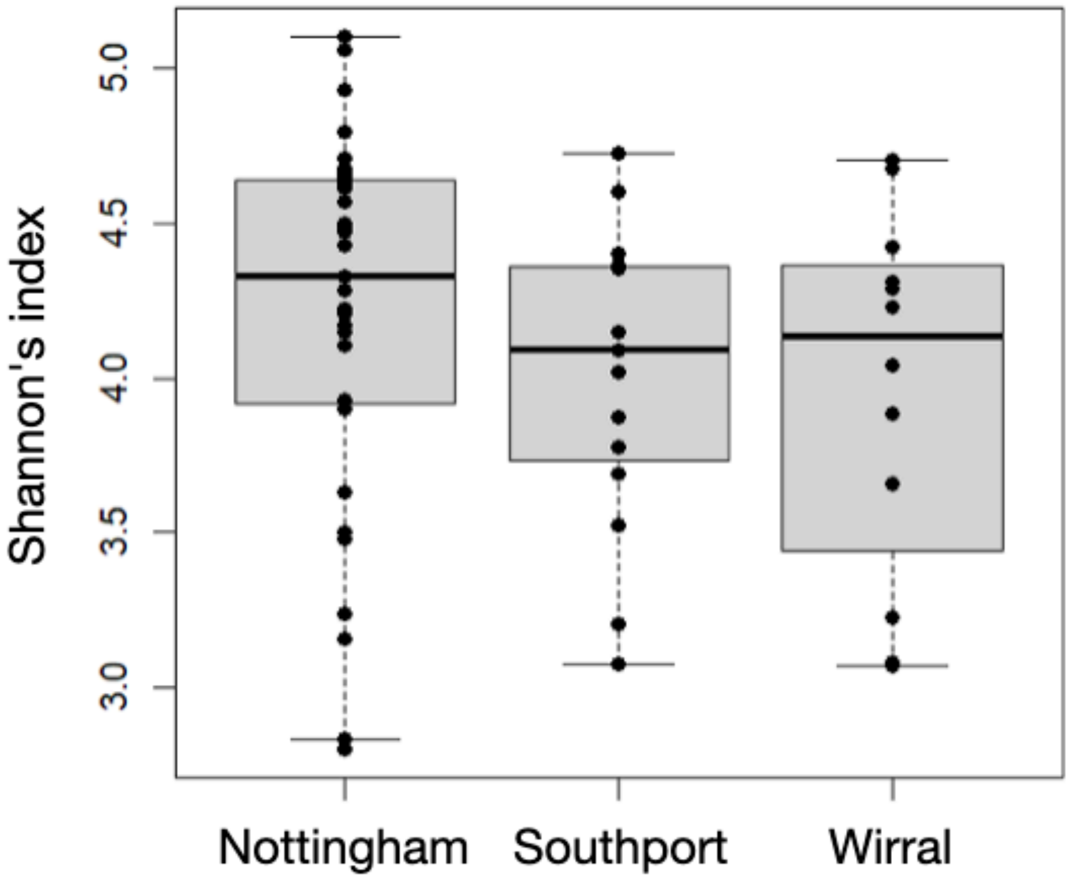

**Supplementary Information 6.** Bacterial microbiome beta diversity calculated using Bray-Curtis dissimilarity from ASV abundance of the pre-sort samples for the three sample sites, which were visualised using a principal coordinate analysis where ellipses are 95% confidence intervals, and which were compared using a permutational multivariate analysis of variance (PERMANOVA) finding that sampling site accounted for 13% of the variation in beta diversity (PERMANOVA: F2 = 4.19, *R*2 = 0.132, *p* = 0.001).

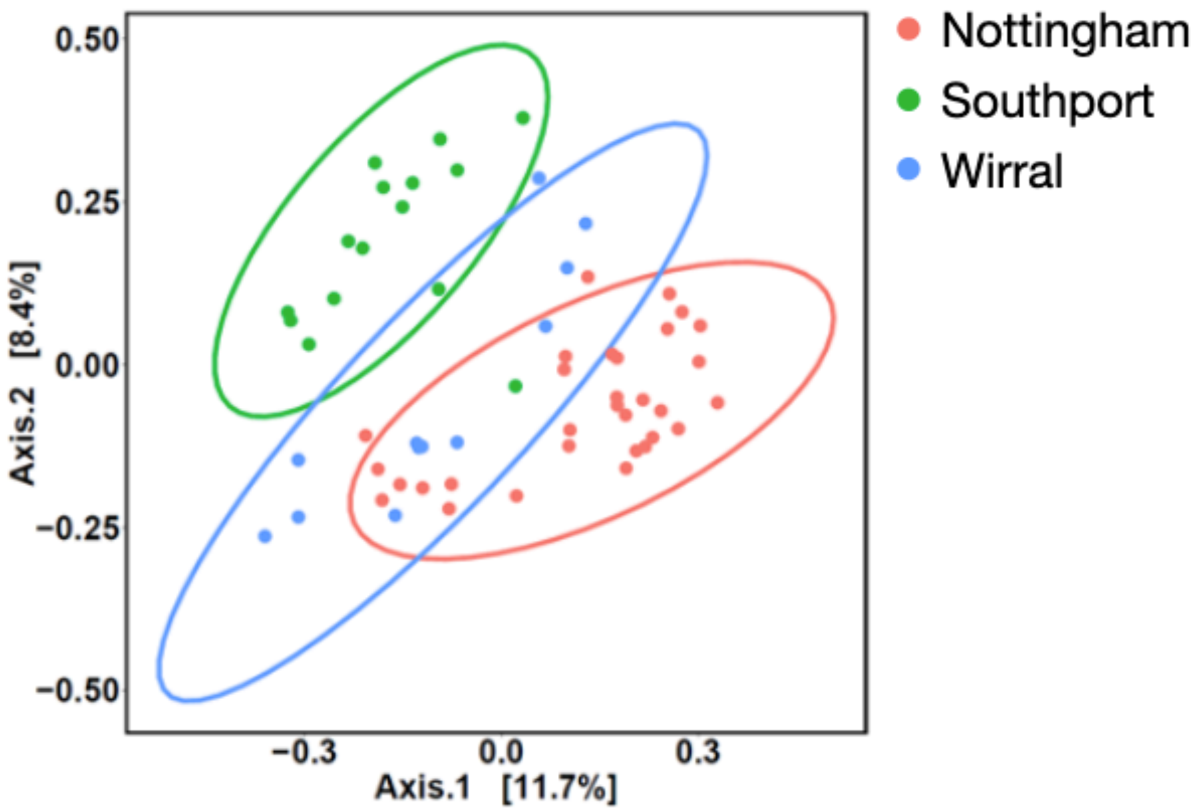

**Supplementary Information 7.** Effect of Bacteroidota relative abundance on the alpha diversity of the bacterial microbiome. Table, parameter estimates of the selected linear model; $R^2c$ is the conditional variance of the model. Figure, the blue line is the model estimate and the grey area the 95% confidence intervals.

| Trait | Best model selected | | Estimates ± SE | t-value | p | $R^2c$ |
|---|---|---|---|---|---|---|
| Shannon index | Bacteroidota relative abundance | Intercept | 4.53 ± 0.12 | 36.52 | < 0.001 | 0.23 |
| | | Bacteroidota | -1.22 ± 0.30 | -4.09 | < 0.001 | |

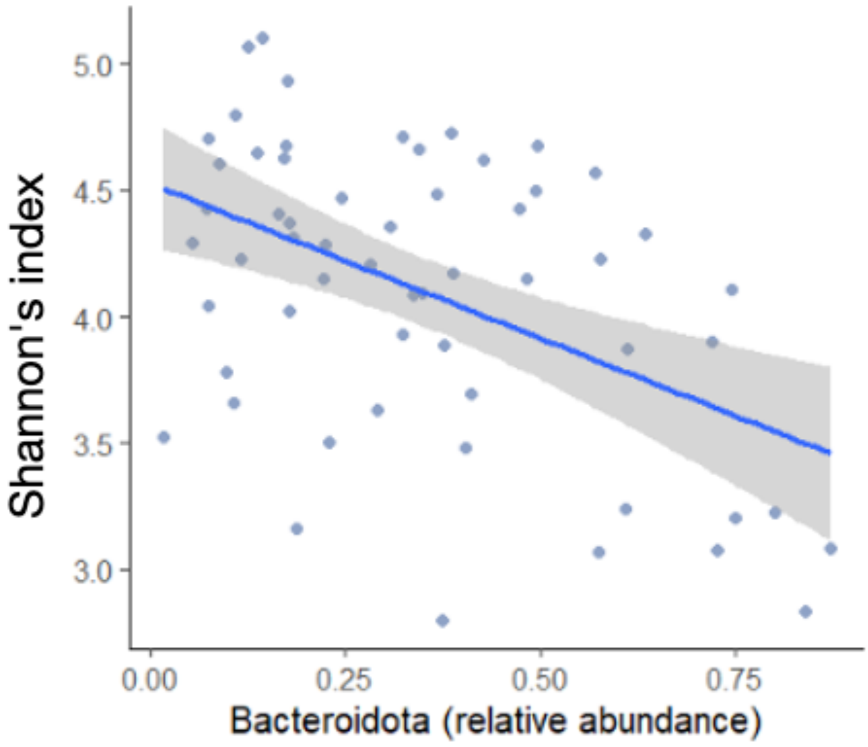

**Supplementary Information 8.** Results of a PERMANOVA analysis of factors affecting beta diversity measured by Bray-Curtis dissimilarity of the bacterial microbiome. In Traits tested: 'F' is faecal, 'C' is Caecal; 'relative abundance ps' is relative abundance of the taxa in the pre-sort fraction; for the 12 ASVs this is relative abundance in the IgA+ fraction. Factors highlighted in green explain ≥5% of the total variation in beta diversity.

| **Trait tested (separately)** | **Effect on beta diversity** | **F2** | **R2** | **p** | **% of variation explained** |
|---|---|---|---|---|---|
| Sample site | Yes | 4.19 | 0.132 | 0.001 | 13 |
| BMI | Yes | 1.84 | 0.032 | 0.007 | 3 |
| Age (weeks) | Yes | 1.42 | 0.024 | 0.038 | 2 |
| Age (< 6 weeks old *vs*. (≥ 6 weeks old) | Yes | 1.76 | 0.030 | 0.007 | 3 |
| Reproductive status | No | 1.34 | 0.023 | 0.069 | 2 |
| [Faecal IgA] | Yes | 1.57 | 0.027 | 0.023 | 3 |
| IgA+ bacteria (%) | Yes | 2.12 | 0.036 | 0.001 | 4 |
| Bacterial Load | Yes | 1.84 | 0.032 | 0.004 | 3 |
| Small Intestine Inflammation | No | 0.79 | 0.014 | 0.842 | 1 |
| Large Intestine Inflammation | No | 1.15 | 0.022 | 0.219 | 2 |
| Caecum Inflammation | No | 1.25 | 0.023 | 0.134 | 2 |
| [Mucin] | No | 1.00 | 0.018 | 0.450 | 2 |
| Apicomplexa (C) (relative abundance) | No | 1.01 | 0.018 | 0.427 | 2 |
| Apicomplexa (F) (relative abundance) | Yes | 1.59 | 0.028 | 0.015 | 3 |
| Ascomycota (C) (relative abundance) | Yes | 3.11 | 0.053 | 0.001 | 5 |
| Ascomycota (F) (relative abundance) | Yes | 1.54 | 0.027 | 0.024 | 3 |
| Basidiomycota (C) (relative abundance) | Yes | 3.67 | 0.062 | 0.001 | 6 |
| Basidiomycota (F) (relative abundance) | Yes | 2.74 | 0.047 | 0.001 | 5 |
| Nematoda (C) (relative abundance) | Yes | 2.81 | 0.048 | 0.001 | 5 |
| Nematoda (F) (relative abundance) | Yes | 1.87 | 0.032 | 0.003 | 3 |
| Eukaryotic (F) alpha diversity (Shannon's) | No | 1.08 | 0.019 | 0.285 | 2 |
| Eukaryotic (C) alpha diversity (Shannon's) | Yes | 1.49 | 0.026 | 0.032 | 3 |
| Bacteriome (F) alpha diversity (Shannon's) | Yes | 4.04 | 0.067 | 0.001 | 7 |
| Firmicutes (relative abundance ps) | Yes | 5.09 | 0.083 | 0.001 | 8 |
| Bacteroidota (relative abundance ps) | Yes | 5.07 | 0.083 | 0.001 | 8 |
| Actinobacteriota (relative abundance ps) | Yes | 1.93 | 0.033 | 0.001 | 3 |
| Proteobacteria (relative abundance ps) | No | 1.22 | 0.021 | 0.129 | 2 |
| ASV24 | No | 1.34 | 0.023 | 0.053 | 2 |
| ASV33 | No | 0.83 | 0.015 | 0.854 | 1 |
| ASV52 | No | 0.85 | 0.015 | 0.828 | 1 |
| ASV57 | No | 0.88 | 0.015 | 0.773 | 1 |
| ASV71 | No | 1.24 | 0.022 | 0.128 | 2 |
| ASV73 | No | 0.84 | 0.015 | 0.778 | 1 |
| ASV135 | No | 0.90 | 0.016 | 0.676 | 2 |
| ASV153 | No | 1.10 | 0.019 | 0.270 | 2 |
| ASV163 | No | 0.76 | 0.013 | 0.903 | 1 |
| ASV168 | No | 1.10 | 0.019 | 0.264 | 2 |
| ASV334 | No | 0.73 | 0.013 | 0.979 | 1 |
| ASV353 | No | 0.85 | 0.015 | 0.773 | 1 |
| ASV 24 + 33 + 52 + 57 + 71 + 73 + 135 + 153 + 163 + 168 + 334 + 353 | No | 1.02 | 0.213 | 0.404 | 21 |

**Supplementary Information 9.** Alpha diversity calculated using Shannon's index from ASV abundance of the pre-sort, IgA$^+$ and IgA$^-$ fractions of the bacterial microbiome, which were compared using Wilcoxon signed-rank tests, finding that they do not differ, Kruskal-Wallis: $\chi^2_2 = 3.95$, $p > 0.05$. Whiskers show the minimum to maximum values in the dataset.

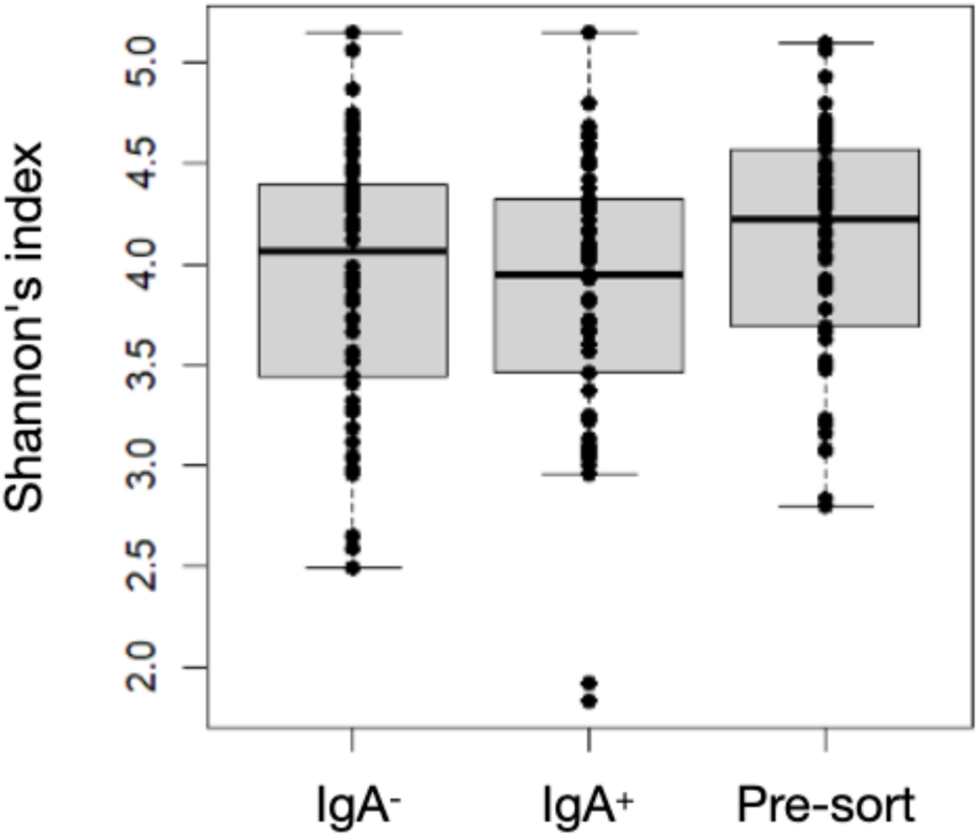

**Supplementary Information 10.** A Principal Component Analysis of Bray-Curtis dissimilarity of the bacterial microbiome where Axis 1 accounts for 10.3% of the variance, and Axis 2 7.7%. Samples are from the Nottingham (N, red), Southport (S, green), and Wirral (W, blue) with the numbers showing individual animal IDs, with fractions (pre-sort, $IgA^+$ and $IgA^-$) shown by symbol shape. Ellipses are 95% confidence intervals.

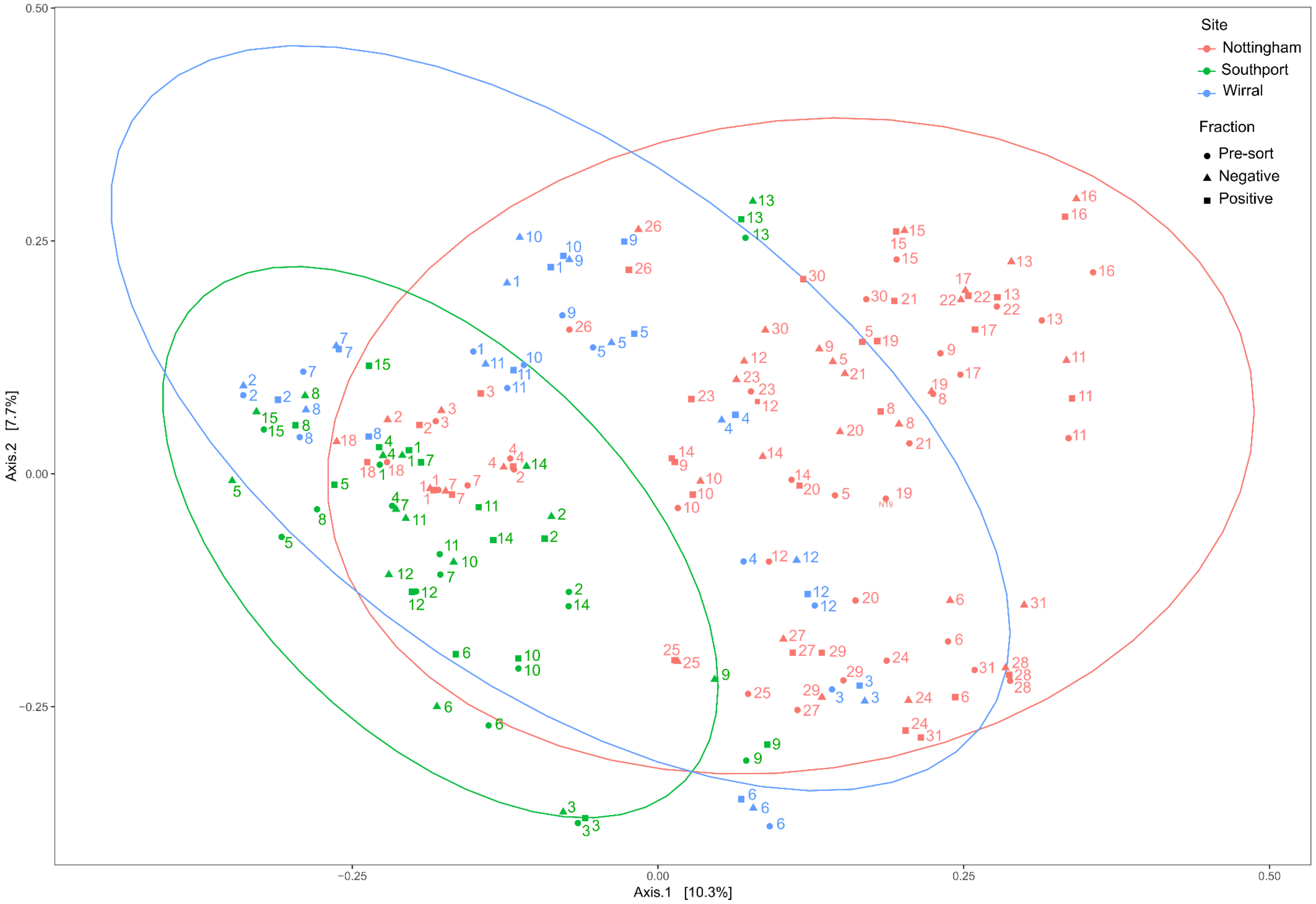

**Supplementary Information 11.** The ranked proportion of the 58 mice that gave a positive IgA probability to an ASV that they harboured.

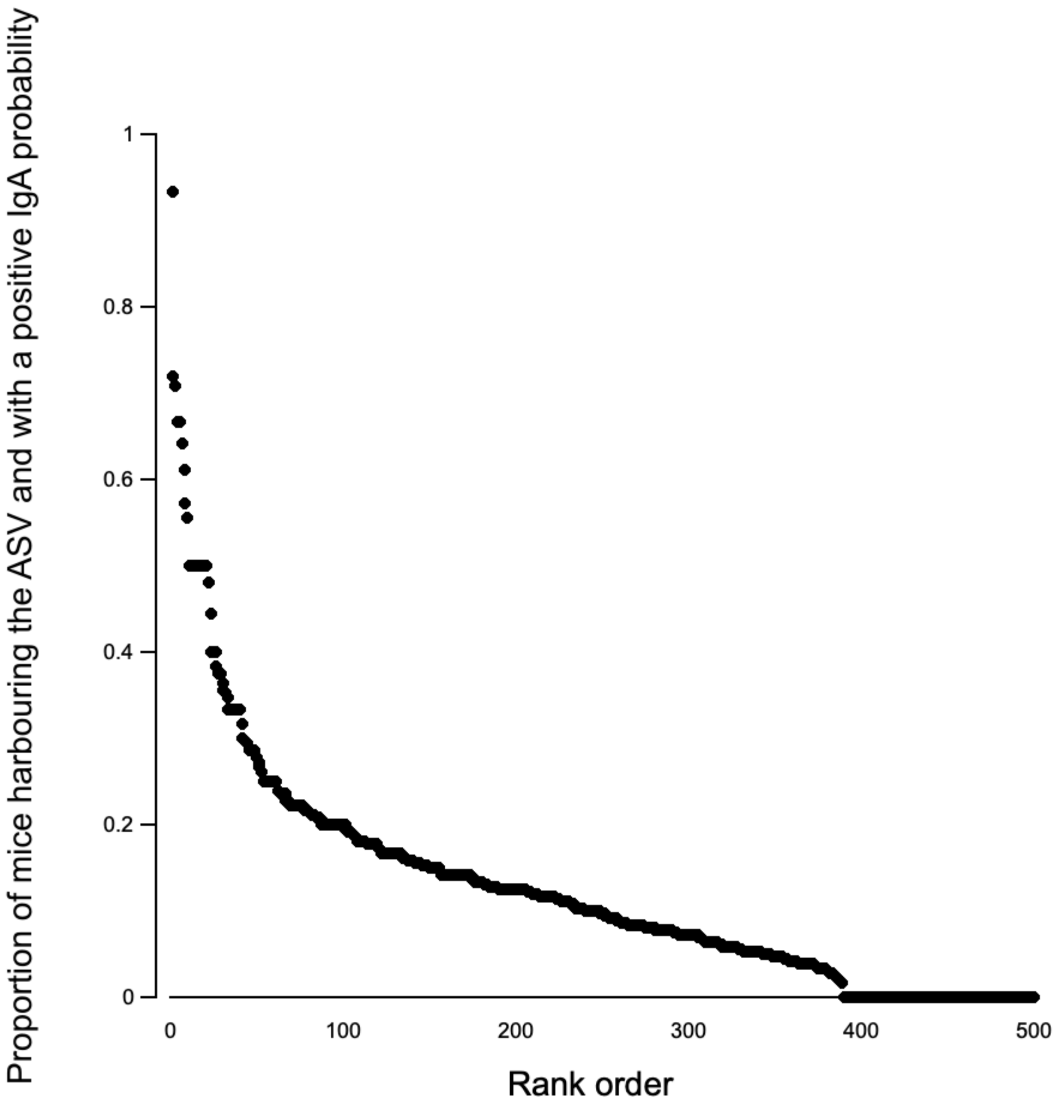

**Supplementary Information 12.** A PCA of traits likely to be relevant to the composition of the gut microbiome (as main text Figure 1) showing 16 of the 19 mice that account for 49% of all ASVs with an IgA$^{+}$ probability. The 16 mice are N02, N05, N07, N13, N14, N17, N21, N29, N30, S01, S02, S06, S07, W04, W06, W10; N06, N18, S06, S13 are not included because they have some missing data that prevents their inclusion in the PCA analysis.

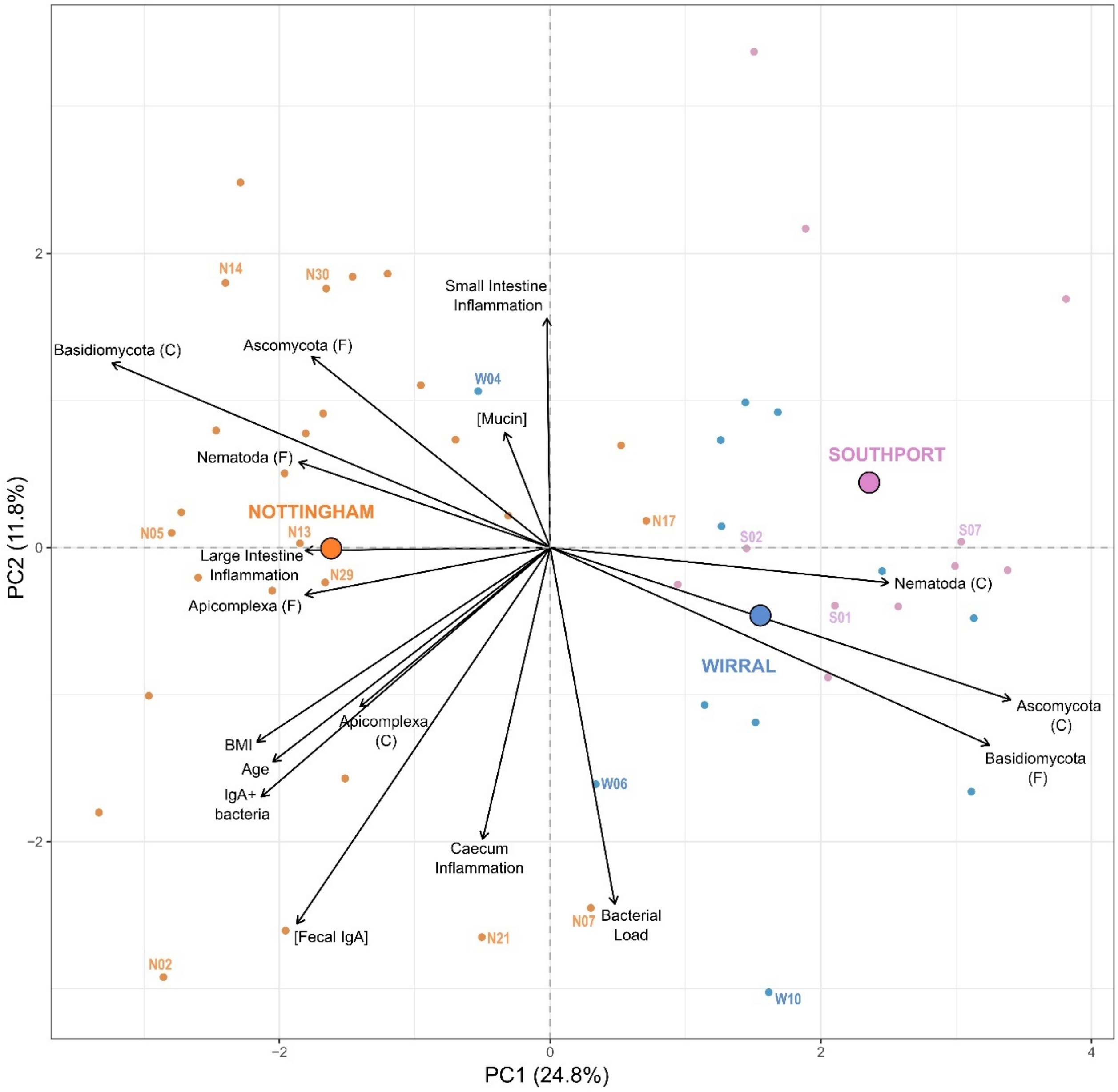

**Supplementary Information 13.** The number of positive IgA probability ASVs harboured by a mouse, against (A) mouse age in weeks, (B) [faecal IgA] in mg IgA / g faeces, (C), the percent of bacteria that are IgA$^{+}$, (D) BMI.

**A**

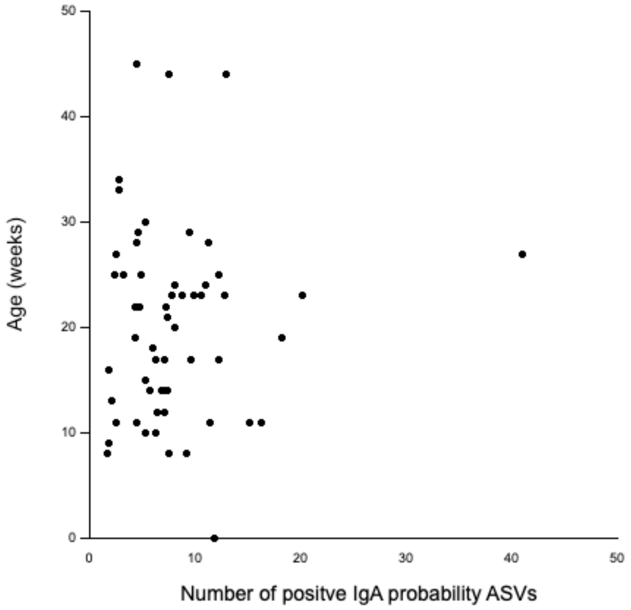

**B**

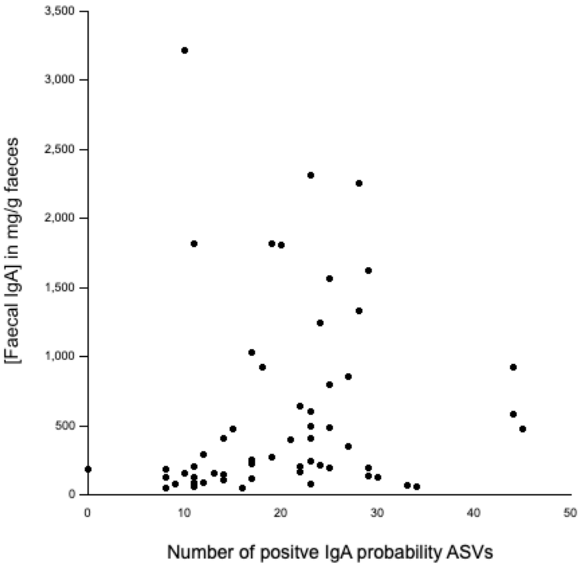

**C**

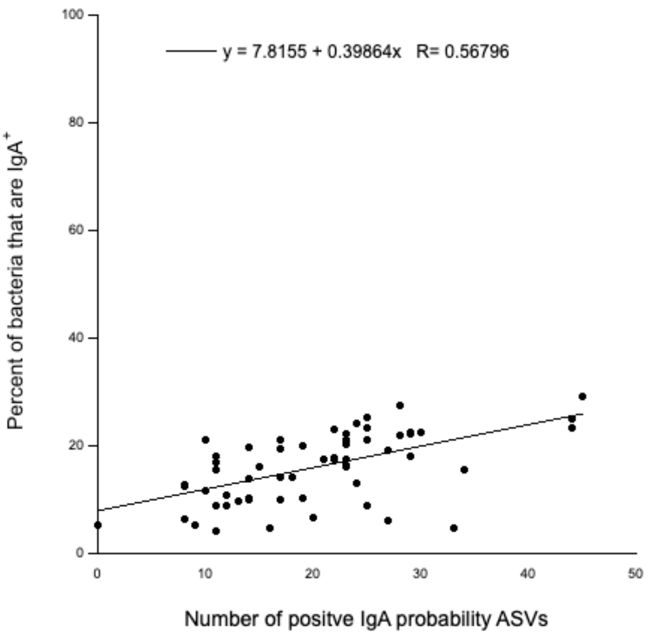

**D**

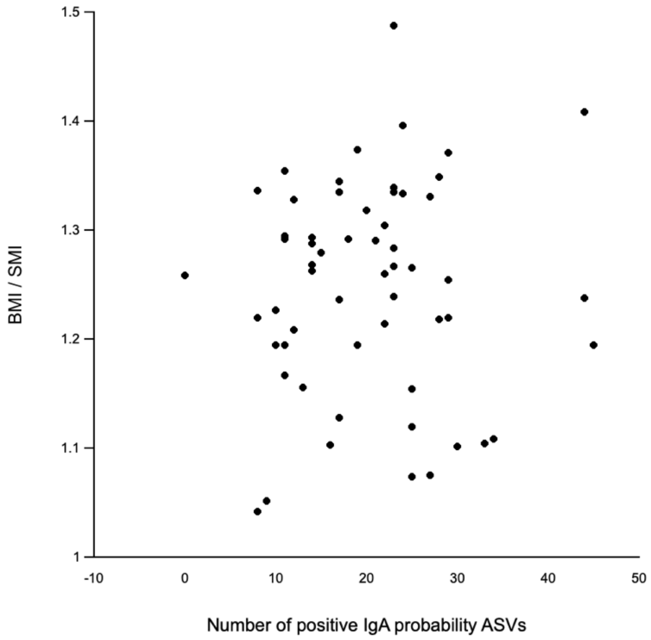

**Supplementary Information 14.** The correlation in abundance in the $IgA^{+}$ fraction of 12 ASVs (from Table 1) highlighted in yellow, and 12 other randomly selected ASVs, together with components of the eukaryotic microbiome. The values in the cells are correlations, with positive values coloured in blue and negative values in red.

SI 14 is an Excel file.

**Supplementary Information 15.** Phyla identified from the faecal and caecal eukaryotic microbiome of wild mice. Phyla are ordered by their relative abundance across all 58 wild mice. Only the phyla with a relative abundance of > 0.1% and > 0.05% for the faecal and caecal microbiome, respectively, are shown. Abundance is the number of sequence reads assigned to each phylum. Prevalence is the percentage of mice from which that phylum was identified. Phyla in bold are known gut residents. N'ham = Nottingham, S'port = Southport.

| | **Phylum** | **Abundance** | | **Prevalence (%)** | | | |
|---|---|---|---|---|---|---|---|
| | | **Relative (%)** | **Total** | **Combined (58)** | **N'ham (31)** | **S'port (15)** | **Wirral (12)** |
| **Faecal** | **Ascomycota** | 42.5 | 1,080,915 | 100 | 100 | 100 | 100 |
| | **Basidiomycota** | 32.0 | 815,080 | 100 | 100 | 100 | 100 |
| | Mucoromycota | 13.4 | 341,999 | 100 | 100 | 100 | 100 |
| | **Apicomplexa** | 9.2 | 233,732 | 88 | 100 | 80 | 67 |
| | **Nematoda** | 0.7 | 18,520 | 60 | 94 | 0 | 50 |
| | Cercozoa | 0.6 | 15,977 | 88 | 94 | 87 | 75 |
| | **Bigyra** | 0.5 | 13,671 | 14 | 19 | 0 | 17 |
| | **Ciliophora** | 0.3 | 6,749 | 78 | 84 | 67 | 75 |
| | Rotifera | 0.1 | 3,613 | 17 | 26 | 13 | 0 |
| | Ochrophyta | 0.1 | 3,585 | 67 | 68 | 67 | 67 |
| | Unclassified | 0.1 | 3,426 | 81 | 87 | 67 | 83 |
| **Caecal** | **Nematoda** | 41.1 | 1,011,293 | 72 | 100 | 27 | 58 |
| | **Basidiomycota** | 24.5 | 603,440 | 100 | 100 | 100 | 100 |
| | **Ascomycota** | 24.2 | 595,891 | 100 | 100 | 100 | 100 |
| | **Apicomplexa** | 6.0 | 147,763 | 88 | 100 | 73 | 75 |
| | Mucoromycota | 3.5 | 86,044 | 98 | 100 | 93 | 100 |
| | **Bigyra** | 0.14 | 3,461 | 7 | 13 | 0 | 0 |
| | Cercozoa | 0.14 | 3,355 | 57 | 52 | 80 | 42 |
| | **Ciliophora** | 0.07 | 1,811 | 52 | 52 | 33 | 75 |
| | Unclassified | 0.07 | 1,680 | 48 | 32 | 67 | 67 |
| | Ochrophyta | 0.06 | 1,358 | 40 | 35 | 33 | 58 |

**Supplementary Information 16.** Relative abundance of eukaryotic phyla in the faecal and caecal eukaryotic microbiome. Relative abundance is the percentage of reads assigned to each phylum. Only the phyla with a sequence abundance of ≥ 200 in ≥ 5% of all mice are shown. Phyla are ordered by their relative abundance across all mice.

| | Phylum | Relative Abundance (%) | | |
|---|---|---|---|---|
| | | Nottingham | Southport | Wirral |
| Faecal | Ascomycota | 46.9 | 42.1 | 34.2 |
| | Basidiomycota | 17.3 | 43.7 | 47.6 |
| | Mucoromycota | 12.0 | 15.0 | 17.0 |
| | Apicomplexa | 19.2 | 0.1 | 0.1 |
| | Nematoda | 1.5 | NA | 0.1 |
| | Cercozoa | 0.9 | 0.5 | 0.1 |
| | Bigyra | 1.1 | NA | < 0.1 |
| | Ciliophora | 0.2 | 0.3 | 0.3 |
| | Ochrophyta | 0.1 | 0.1 | 0.3 |
| Caecal | Nematoda | 64.9 | <0.1 | 7.4 |
| | Ascomycota | 16.3 | 38.4 | 35.0 |
| | Basidiomycota | 6.1 | 60.3 | 45.8 |
| | Apicomplexa | 9.7 | 0.1 | 0.2 |
| | Mucoromycota | 2.3 | 1.0 | 10.9 |
| | Ciliophora | 0.1 | <0.1 | 0.2 |

**Supplementary Information 17.** Results of an analysis of factors affecting alpha diversity (Shannon's index) of the (A) caecal and (B) faecal eukaryotic microbiome. Stepwise linear models were constructed retaining significant factors at each step, testing in sequence for (i) mouse traits (site, sex, age, BMI, reproductive status), (ii) immune-related traits ([faecal IgA], bacterial load, proportion $IgA^+$ bacteria), (iii) gut inflammation ([mucin], small intestine, large intestine and caecum inflammation), (iv) the faecal eukaryotic (relative abundance of Apicomplexa, Basidiomycota, Nematoda, Ascomycota), (v) the caecal eukaryotic (relative abundance of Apicomplexa, Basidiomycota, Nematoda, Ascomycota) and (vi) the main bacterial phyla (relative abundance of Firmicutes, Bacteroidota, Actinobacteriota and Proteobacteria). F is Faecal; C is Caecal; SE is standard error; * = p < .05, ** = p < .01 and *** = p < .001; $R^2c$ is the conditional variance of the model.

**A**

| Trait | Best model selected | | Estimates ± SE | t-value | p | $R^2c$ |
|---|---|---|---|---|---|---|
| Shannon index eukaryotic (C) | Small Intestine Inflammation + Bacterial Load + Nematoda (F) + Basidiomycota (C) + Apicomplexa (C) + Ascomycota (C) + Nematoda (C) | Intercept | 6.04 ± 0.82 | 7.35 | *** | 0.66 |
| | | Small Intestine Inflammation | -0.20 ± 0.05 | -3.63 | *** | |
| | | Bacterial Load | $1.11 \times 10^{-9} \pm 1.49 \times 10^{-9}$ | 0.74 | - | |
| | | Nematoda (F) | -0.72 ± 2.15 | -0.34 | - | |
| | | Basidiomycota (C) | -4.83 ± 0.82 | -5.90 | *** | |
| | | Apicomplexa (C) | -4.20 ± 0.85 | -4.92 | *** | |
| | | Ascomycota (C) | -4.95 ± 0.87 | -5.71 | *** | |
| | | Nematoda (C) | -3.20 ± 0.86 | -3.72 | *** | |

**B**

| Trait | Best model selected | | Estimates ± SE | t-value | p | $R^2c$ |
|---|---|---|---|---|---|---|
| Shannon index eukaryotic (F) | Small Intestine Inflammation + Caecal Inflammation + Apicomplexa (F) + Basidiomycota (F) + Apicomplexa (C) + Ascomycota (C) + Actinobacteriota | Intercept | 4.17 ± 0.40 | 10.31 | *** | 0.56 |
| | | Small Intestine Inflammation | -0.17 ± 0.05 | -3.17 | ** | |
| | | Caecal Inflammation | -0.24 ± 0.15 | -1.60 | - | |
| | | Apicomplexa (F) | -1.22 ± 0.47 | -2.60 | * | |
| | | Basidiomycota (F) | -1.36 ± 0.49 | -2.80 | ** | |
| | | Apicomplexa (C) | -0.83 ± 0.39 | -2.12 | * | |
| | | Ascomycota (C) | -1.01 ± 0.45 | -2.26 | * | |
| | | Actinobacteriota | -3.54 ± 1.74 | -2.03 | * | |

**Supplementary Information 18.** Results of a PERMANOVA analysis of factors affecting beta diversity (Bray-Curtis dissimilarity) of the (A) caecal and (B) faecal eukaryotic microbiome. In Traits tested: 'F' is faecal, 'C' is Caecal; 'relative abundance ps' is relative abundance of the taxa in the pre-sort fraction; for the 12 ASVs this is relative abundance in the IgA+ fraction. Factors highlighted in green explain ≥5% of the total variation in beta diversity.

**A**

| Trait tested (separately) | Effect on beta diversity | F2 | R2 | p | % of variation explained |
|---|---|---|---|---|---|
| Sample site | Yes | 3.96 | 0.126 | 0.001 | 13 |
| BMI | Yes | 2.77 | 0.047 | 0.006 | 5 |
| Age (weeks) | No | 1.01 | 0.018 | 0.455 | 2 |
| Age (< 6 weeks old *vs.* (≥ 6 weeks old) | Yes | 2.02 | 0.035 | 0.032 | 4 |
| Reproductive status | No | 1.50 | 0.026 | 0.138 | 3 |
| [Faecal IgA] | No | 1.38 | 0.024 | 0.170 | 2 |
| IgA+ bacteria (%) | Yes | 2.02 | 0.035 | 0.035 | 4 |
| Bacterial Load | No | 1.45 | 0.025 | 0.170 | 3 |
| Small Intestine Inflammation | No | 0.93 | 0.017 | 0.505 | 2 |
| Large Intestine Inflammation | No | 1.22 | 0.023 | 0.261 | 2 |
| Caecum Inflammation | No | 1.23 | 0.023 | 0.270 | 2 |
| [Mucin] | No | 1.02 | 0.019 | 0.410 | 2 |
| Apicomplexa (C) (relative abundance) | Yes | 1.92 | 0.033 | 0.042 | 3 |
| Apicomplexa (F) (relative abundance) | Yes | 2.61 | 0.045 | 0.013 | 5 |
| Ascomycota (C) (relative abundance) | Yes | 8.73 | 0.134 | 0.001 | 13 |
| Ascomycota (F) (relative abundance) | Yes | 3.55 | 0.060 | 0.002 | 6 |
| Basidiomycota (C) (relative abundance) | Yes | 9.70 | 0.148 | 0.001 | 15 |
| Basidiomycota (F) (relative abundance) | Yes | 7.43 | 0.117 | 0.001 | 12 |
| Nematoda (C) (relative abundance) | Yes | 4.83 | 0.079 | 0.001 | 8 |
| Nematoda (F) (relative abundance) | Yes | 3.20 | 0.054 | 0.003 | 5 |
| Eukaryotic (F) alpha diversity (Shannon's) | Yes | 5.28 | 0.086 | 0.001 | 9 |
| Eukaryotic (C) alpha diversity (Shannon's) | Yes | 4.19 | 0.070 | 0.001 | 7 |
| Bacteriome (F) alpha diversity (Shannon's) | No | 0.58 | 0.010 | 0.825 | 1 |
| Firmicutes (relative abundance ps) | No | 1.06 | 0.019 | 0.385 | 2 |
| Bacteroidota | No | 1.37 | 0.024 | 0.173 | 2 |
| Actinobacteriota | Yes | 1.87 | 0.032 | 0.049 | 3 |
| Proteobacteria | Yes | 2.69 | 0.046 | 0.006 | 5 |
| ASV24 (rel. ab. in positive fraction) | No | 0.58 | 0.010 | 0.819 | 1 |
| ASV33 | No | 0.68 | 0.012 | 0.728 | 1 |
| ASV52 | No | 0.70 | 0.012 | 0.714 | 1 |
| ASV57 | No | 0.69 | 0.012 | 0.742 | 1 |
| ASV71 | No | 0.92 | 0.014 | 0.584 | 1 |
| ASV73 | No | 1.62 | 0.028 | 0.091 | 3 |
| ASV135 | No | 1.48 | 0.026 | 0.157 | 3 |
| ASV153 | No | 1.71 | 0.030 | 0.083 | 3 |
| ASV163 | No | 1.21 | 0.021 | 0.248 | 2 |
| ASV168 | No | 1.36 | 0.024 | 0.190 | 2 |
| ASV334 | No | 0.32 | 0.006 | 0.999 | 1 |
| ASV353 | No | 0.90 | 0.016 | 0.532 | 1 |
| ASV 24 + 33 + 52 + 57 + 71 + 73 + 135 + 153 + 163 + 168 + 334 + 353 | No | 1.02 | 0.214 | 0.376 | 21 |

**B**

| Trait tested (separately) | Effect on beta diversity | F2 | R2 | p | % of variation explained |
|---|---|---|---|---|---|
| Sample site | Yes | 8.05 | 0.227 | 0.001 | 23 |
| BMI | Yes | 2.44 | 0.042 | 0.021 | 4 |
| Age (weeks) | Yes | 2.28 | 0.039 | 0.018 | 4 |
| Age (< 6 weeks old *vs.* (≥ 6 weeks old) | No | 1.23 | 0.022 | 0.237 | 2 |
| Reproductive status | No | 1.82 | 0.031 | 0.060 | 3 |
| [Faecal IgA] | Yes | 3.57 | 0.060 | 0.005 | 6 |
| IgA+ bacteria (%) | Yes | 2.40 | 0.041 | 0.020 | 4 |
| Bacterial Load | No | 1.10 | 0.010 | 0.353 | 1 |
| Small Intestine Inflammation | No | 0.85 | 0.015 | 0.573 | 2 |
| Large Intestine Inflammation | No | 1.42 | 0.027 | 0.157 | 2 |
| Caecum Inflammation | No | 1.34 | 0.025 | 0.199 | 3 |
| [Mucin] | No | 0.82 | 0.015 | 0.589 | 2 |
| Apicomplexa (C) (relative abundance) | Yes | 1.93 | 0.033 | 0.041 | 3 |
| Apicomplexa (F) (relative abundance) | Yes | 6.10 | 0.098 | 0.001 | 10 |
| Ascomycota (C) (relative abundance) | Yes | 12.7 | 0.185 | 0.001 | 19 |
| Ascomycota (F) (relative abundance) | Yes | 5.77 | 0.093 | 0.001 | 9 |
| Basidiomycota (C) (relative abundance) | Yes | 9.18 | 0.141 | 0.001 | 14 |
| Basidiomycota (F) (relative abundance) | Yes | 16.7 | 0.230 | 0.001 | 23 |
| Nematoda (C) (relative abundance) | Yes | 3.99 | 0.067 | 0.002 | 7 |
| Nematoda (F) (relative abundance) | Yes | 3.46 | 0.058 | 0.002 | 6 |
| Eukaryotic (F) alpha diversity (Shannon's) | Yes | 8.45 | 0.131 | 0.001 | 13 |
| Eukaryotic (C) alpha diversity (Shannon's) | No | 1.33 | 0.023 | 0.149 | 2 |
| Bacteriome (F) alpha diversity (Shannon's) | No | 0.77 | 0.014 | 0.655 | 1 |
| Firmicutes (relative abundance ps) | No | 1.03 | 0.018 | 0.385 | 2 |
| Bacteroidota | No | 1.25 | 0.022 | 0.231 | 2 |
| Actinobacteriota | Yes | 1.90 | 0.033 | 0.046 | 5 |
| Proteobacteria | Yes | 2.24 | 0.039 | 0.039 | 4 |
| ASV24 | No | 0.74 | 0.013 | 0.687 | 1 |
| ASV33 | No | 1.60 | 0.028 | 0.086 | 3 |
| ASV52 | No | 1.76 | 0.031 | 0.059 | 1 |
| ASV57 | No | 1.72 | 0.030 | 0.064 | 3 |
| ASV71 | No | 0.59 | 0.010 | 0.589 | 1 |
| ASV73 | No | 0.74 | 0.013 | 0.691 | 1 |
| ASV135 | No | 1.18 | 0.021 | 0.297 | 2 |
| ASV153 | No | 1.23 | 0.021 | 0.258 | 2 |
| ASV163 | No | 1.28 | 0.022 | 0.224 | 2 |
| ASV168 | No | 2.27 | 0.039 | 0.019 | 4 |
| ASV334 | No | 0.63 | 0.011 | 0.860 | 1 |
| ASV353 | No | 1.62 | 0.028 | 0.075 | 3 |
| ASV 24 + 33 + 52 + 57 + 71 + 73 + 135 + 153 + 163 + 168 + 334 + 353 | No | 0.97 | 0.206 | 0.549 | 21 |
| 12 random ASVs | No | 1.01 | 0.211 | 0.447 | 21 |

**Supplementary Information 19.** For alpha diversity of (A) caecal eukaryotic and (B) faecal eukaryotic models as **Supplementary Information 17** but were Apicomplexa is replaced by the relative abundance of *Eimeria* (shown in bold, below, compared with **Supplementary Information 17**), showing that the model results remain essentially unchanged; (C) a PERMANOVA analysis testing the effect of *Eimeria* relative abundance on beta diversity (Bray-Curtis dissimilarity). F is Faecal; C is Caecal; SE is standard error; * = p < .05, ** = p < .01 and *** = p < .001; $R^2c$ is the conditional variance of the model.

**A**

| Trait | Best model selected | | Estimates ± SE | t-value | p | $R^2c$ |
|---|---|---|---|---|---|---|
| Shannon index eukaryotic (C) | Small Intestine Inflammation + Bacterial Load + Nematoda (F) + Basidiomycota (C) + ***Eimeria*** (C) + Ascomycota (C) + Nematoda (C) | Intercept | 6.11 ± 0.79 | 7.69 | *** | 0.67 |
| | | Small Intestine Inflammation | -0.19 ± 0.05 | -3.65 | *** | |
| | | Bacterial Load | $1.19 \times 10^{-9} \pm 1.46 \times 10^{-9}$ | 0.81 | - | |
| | | Nematoda (F) | -0.69 ± 2.10 | -0.32 | - | |
| | | Basidiomycota (C) | -4.92 ± 0.83 | -6.18 | *** | |
| | | ***Eimeria*** (C) | -4.31 ± 0.83 | -5.20 | *** | |
| | | Ascomycota (C) | -5.03 ± 0.84 | -5.98 | *** | |
| | | Nematoda (C) | -3.30 ± 0.84 | -3.93 | *** | |

**B**

| Trait | Best model selected | | Estimates ± SE | t-value | p | $R^2c$ |
|---|---|---|---|---|---|---|
| Shannon index eukaryotic (F) | Small Intestine Inflammation + Caecal Inflammation + Apicomplexa (F) + Basidiomycota (F) + ***Eimeria*** (C) + Ascomycota (C) + Actinobacteriota | Intercept | 4.16 ± 0.40 | 10.29 | *** | 0.56 |
| | | Small Intestine Inflammation | -0.17 ± 0.05 | -3.14 | ** | |
| | | Caecal Inflammation | -0.24 ± 0.15 | -1.63 | - | |
| | | Apicomplexa (F) | -1.21 ± 0.47 | -2.58 | * | |
| | | Basidiomycota (F) | -1.34 ± 0.49 | -2.77 | ** | |
| | | ***Eimeria*** (C) | -0.82 ± 0.39 | -2.25 | * | |
| | | Ascomycota (C) | -1.01 ± 0.45 | -2.25 | * | |
| | | Actinobacteriota | -3.47 ± 1.74 | -2.00 | * | |

**C**

| Diversity | Trait tested (separately) | Effect on beta diversity | F2 | R2 | P | | % of variation explained |
|---|---|---|---|---|---|---|---|
| Bacterial | *Eimeria* relative abundance (F) | Yes | 1.51 | 0.026 | 0.03 | * | 3 |
| Bacterial | *Eimeria* relative abundance (C) | No | 1.00 | 0.017 | 0.454 | - | 2 |
| Eukaryotic (C) | *Eimeria* relative abundance (F) | Yes | 2.58 | 0.044 | 0.002 | ** | 4 |
| Eukaryotic (C) | *Eimeria* relative abundance (C) | Yes | 5.30 | 0.086 | 0.001 | *** | 9 |
| Eukaryotic (F) | *Eimeria* relative abundance (F) | Yes | 5.83 | 0.094 | 0.001 | *** | 9 |
| Eukaryotic (F) | *Eimeria* relative abundance (C) | No | 1.89 | 0.033 | 0.055 | - | 3 |

**Supplementary Information 20.** Networks of the 75 most abundant ASVs; if any of the 75 ASVs were not associated with other ASVs then they are not shown in the network, to improve clarity. Common coloured nodes are clusters, which are groups of nodes that are highly connected to one another but have a small number of connections to other nodes. The size of nodes shows the node eigenvector centrality. Links between taxa are shown by lines, with positive associations shown in green, and negative associations shown in red; line thickness shows the degree of the positive or negative effects. The taxa names show in parentheses whether they are bacterial (B) or eukaryotic (E). The networks are of,(A) all mice, (B) mice from the Nottingham site, (C) mice from the Southport site, and (D) mice from the Wirral site. Within each network the number of positive and negative bacterial-eukaryotic interactions are 8 and 9, 6 and 4, 9 and 21, 14 and 7, for A, B, C, D, respectively.

A

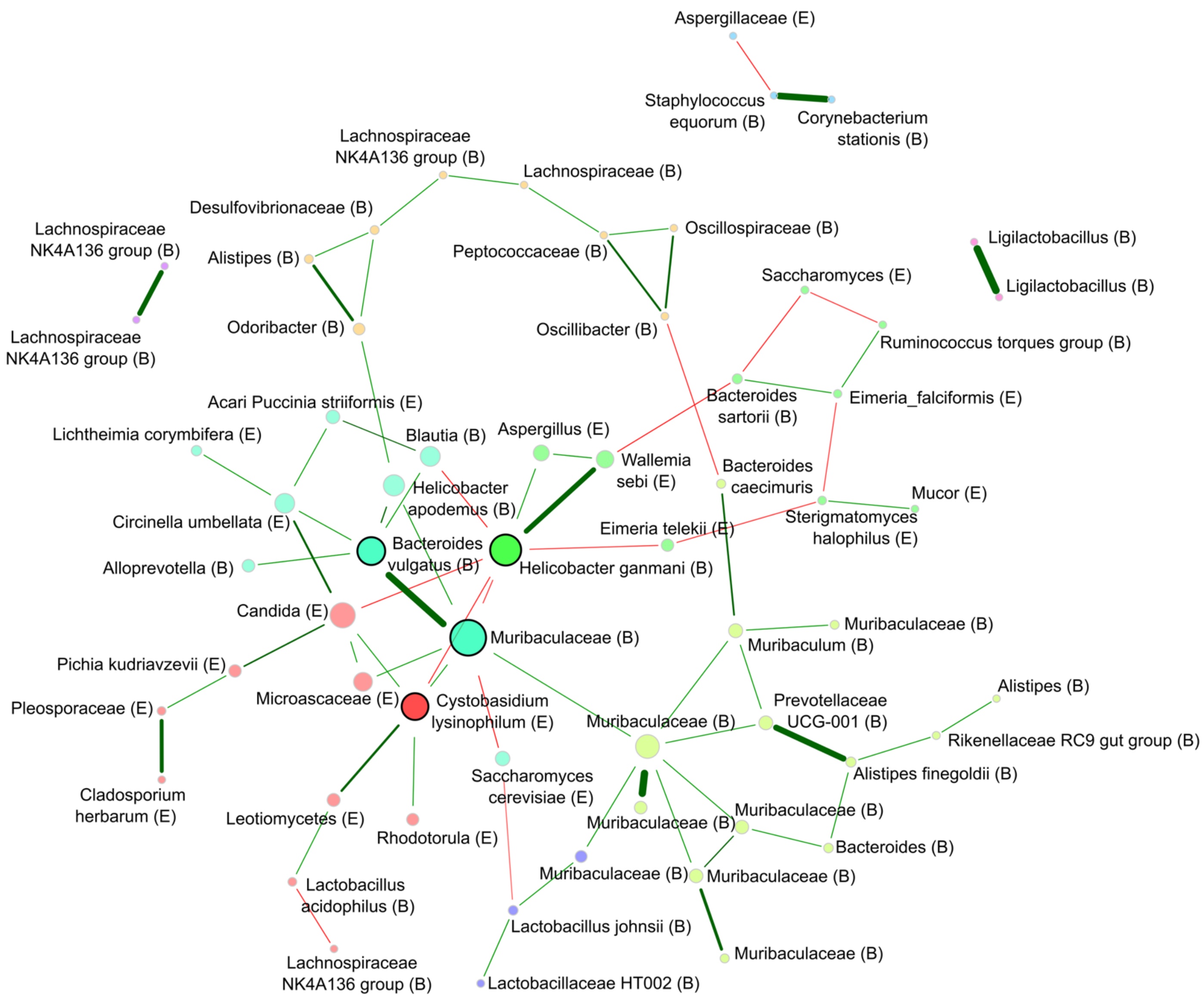
Aspergillaceae (E)
Staphylococcus equorum (B)
Corynebacterium stationis (B)
Lachnospiraceae NK4A136 group (B)
Lachnospiraceae (B)
Desulfovibrionaceae (B)
Oscillospiraceae (B)
Lachnospiraceae NK4A136 group (B)
Alistipes (B)
Peptococcaceae (B)
Ligilactobacillus (B)
Saccharomyces (E)
Ligilactobacillus (B)
Lachnospiraceae NK4A136 group (B)
Odoribacter (B)
Oscillibacter (B)
Ruminococcus torques group (B)
Bacteroides sartorii (B)
Eimeria_falciformis (E)
Acari Puccinia striiformis (E)
Lichtheimia corymbifera (E)
Blautia (B)
Aspergillus (E)
Wallemia sebi (E)
Helicobacter apodemus (B)
Bacteroides caecimuris
Mucor (E)
Sterigmatomyces halophilus (E)
Circinella umbellata (E)
Eimeria telekii (E)
Bacteroides vulgatus (B)
Helicobacter ganmani (B)
Alloprevotella (B)
Candida (E)
Muribaculaceae (B)
Muribaculaceae (B)
Muribaculum (B)
Pichia kudriavzevii (E)
Alistipes (B)
Microascaceae (E)
Cystobasidium lysinophilum (E)
Prevotellaceae UCG-001 (B)
Pleosporaceae (E)
Muribaculaceae (B)
Rikenellaceae RC9 gut group (B)
Alistipes finegoldii (B)
Cladosporium herbarum (E)
Saccharomyces cerevisiae (E)
Muribaculaceae (B)
Leotiomycetes (E)
Rhodotorula (E)
Muribaculaceae (B)
Bacteroides (B)
Muribaculaceae (B)
Muribaculaceae (B)
Lactobacillus acidophilus (B)
Lactobacillus johnsii (B)
Muribaculaceae (B)
Lachnospiraceae NK4A136 group (B)
Lactobacillaceae HT002 (B)

**B**

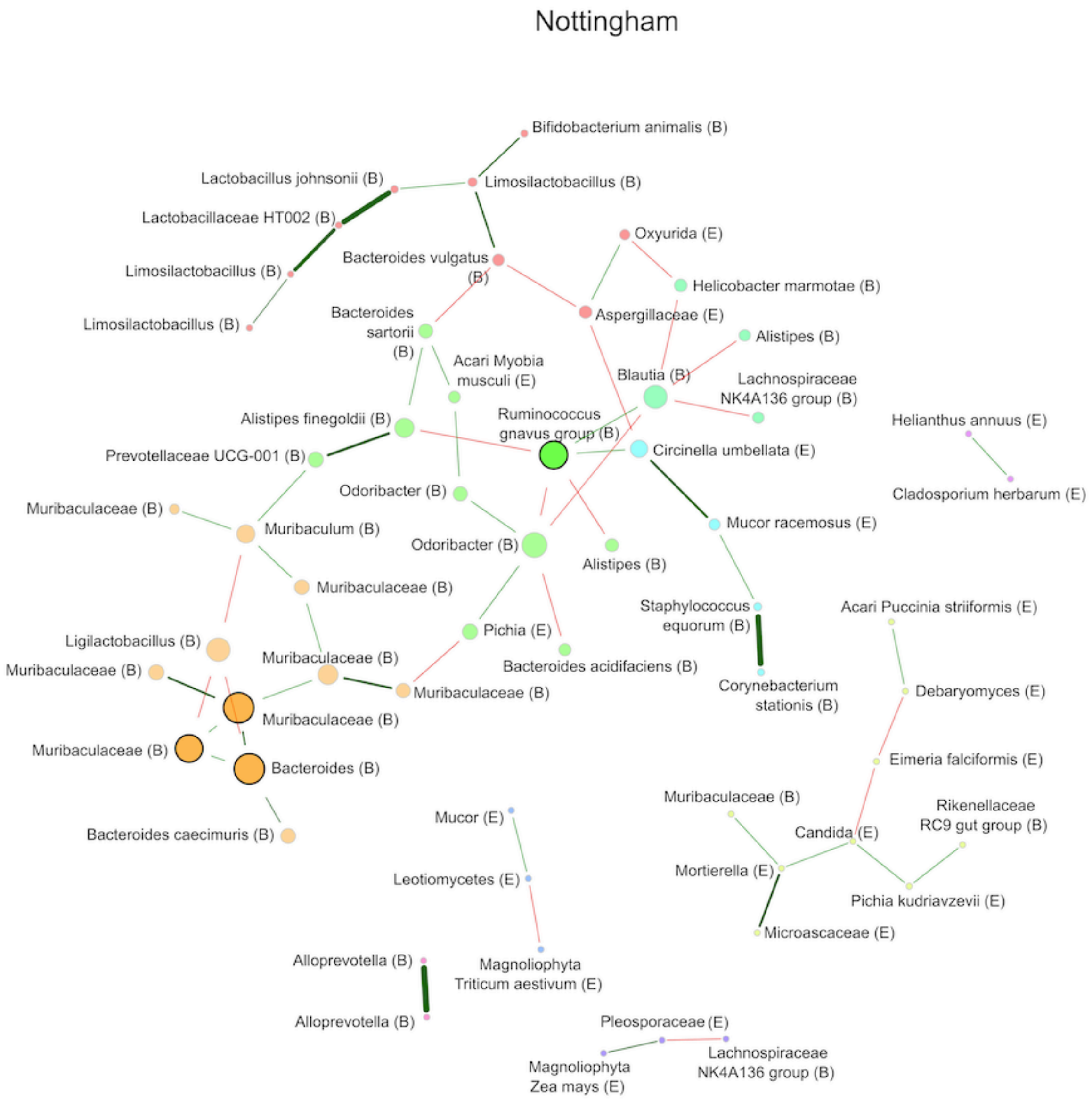
Nottingham
Bifidobacterium animalis (B)
Lactobacillus johnsonii (B)
Limosilactobacillus (B)
Lactobacillaceae HT002 (B)
Oxyurida (E)
Bacteroides vulgatus (B)
Limosilactobacillus (B)
Helicobacter marmotae (B)
Aspergillaceae (E)
Limosilactobacillus (B)
Bacteroides sartorii (B)
Alistipes (B)
Acari Myobia musculi (E)
Blautia (B)
Lachnospiraceae NK4A136 group (B)
Ruminococcus gnavus group (B)
Helianthus annuus (E)
Alistipes finegoldii (B)
Circinella umbellata (E)
Prevotellaceae UCG-001 (B)
Cladosporium herbarum (E)
Odoribacter (B)
Muribaculaceae (B)
Mucor racemosus (E)
Muribaculum (B)
Odoribacter (B)
Alistipes (B)
Muribaculaceae (B)
Staphylococcus equorum (B)
Acari Puccinia striiformis (E)
Pichia (E)
Ligilactobacillus (B)
Muribaculaceae (B)
Bacteroides acidifaciens (B)
Muribaculaceae (B)
Corynebacterium stationis (B)
Debaryomyces (E)
Muribaculaceae (B)
Muribaculaceae (B)
Muribaculaceae (B)
Eimeria falciformis (E)
Bacteroides (B)
Muribaculaceae (B)
Rikenellaceae RC9 gut group (B)
Candida (E)
Mucor (E)
Bacteroides caecimuris (B)
Mortierella (E)
Leotiomycetes (E)
Pichia kudriavzevii (E)
Microascaceae (E)
Alloprevotella (B)
Magnoliophyta Triticum aestivum (E)
Alloprevotella (B)
Pleosporaceae (E)
Lachnospiraceae NK4A136 group (B)
Magnoliophyta Zea mays (E)

**C**

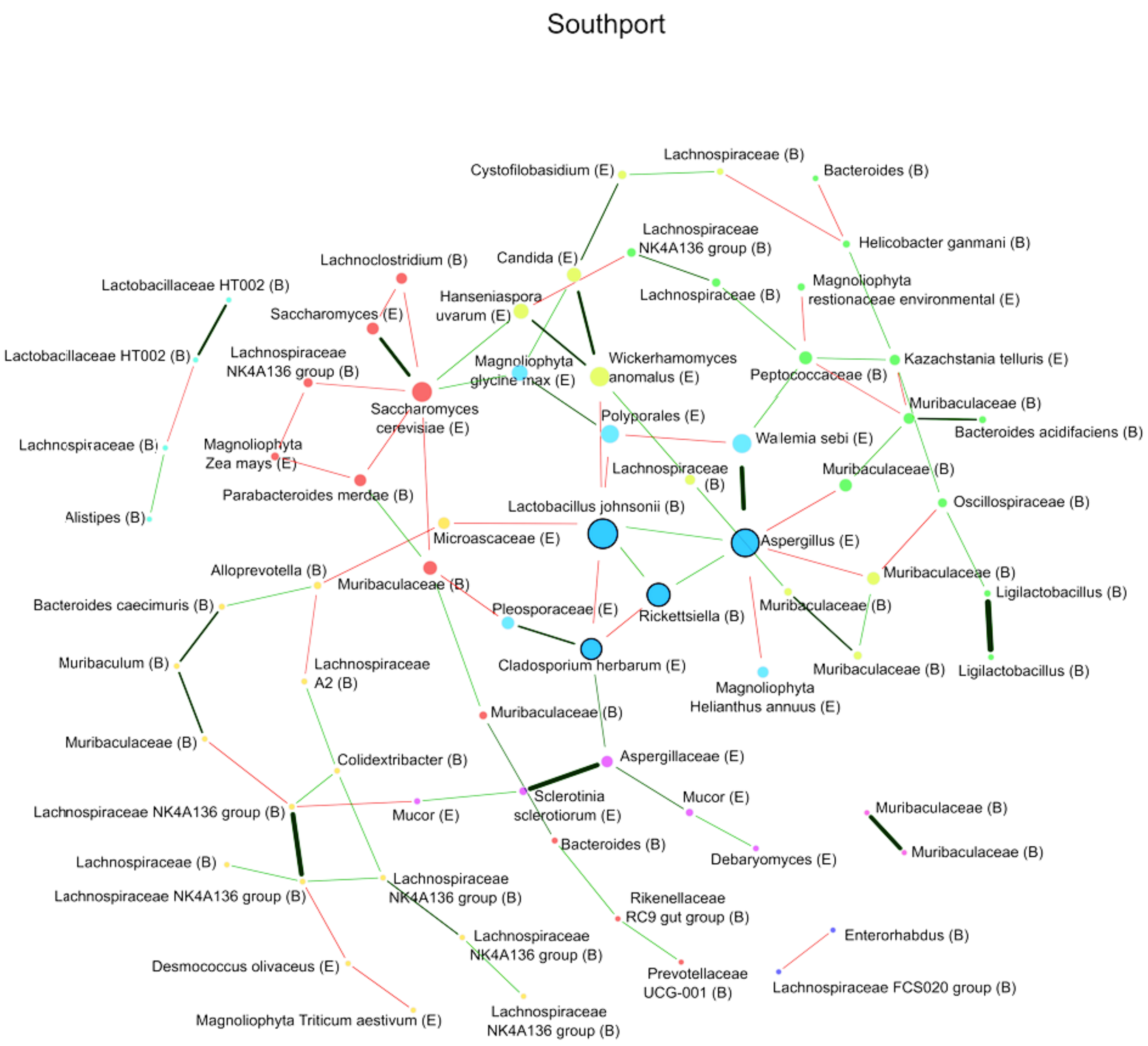
Southport
Cystofilobasidium (E)
Lachnospiraceae (B)
Bacteroides (B)
Lachnospiraceae NK4A136 group (B)
Helicobacter ganmani (B)
Lachnoclostridium (B)
Candida (E)
Lactobacillaceae HT002 (B)
Hanseniaspora uvarum (E)
Lachnospiraceae (B)
Magnoliophyta restionaceae environmental (E)
Saccharomyces (E)
Lactobacillaceae HT002 (B)
Lachnospiraceae NK4A136 group (B)
Magnoliophyta glycine max (E)
Wickerhamomyces anomalus (E)
Peptococcaceae (B)
Kazachstania telluris (E)
Saccharomyces cerevisiae (E)
Muribaculaceae (B)
Bacteroides acidifaciens (B)
Polyporales (E)
Lachnospiraceae (B)
Magnoliophyta Zea mays (E)
Walemia sebi (E)
Lachnospiraceae (B)
Muribaculaceae (B)
Parabacteroides merdae (B)
Alistipes (B)
Lactobacillus johnsonii (B)
Oscillospiraceae (B)
Microascaceae (E)
Aspergillus (E)
Alloprevotella (B)
Muribaculaceae (B)
Muribaculaceae (B)
Ligilactobacillus (B)
Bacteroides caecimuris (B)
Pleosporaceae (E)
Muribaculaceae (B)
Rickettsiella (B)
Muribaculum (B)
Lachnospiraceae A2 (B)
Cladosporium herbarum (E)
Muribaculaceae (B)
Ligilactobacillus (B)
Magnoliophyta Helianthus annuus (E)
Muribaculaceae (B)
Muribaculaceae (B)
Colidextribacter (B)
Aspergillaceae (E)
Mucor (E)
Sclerotinia sclerotiorum (E)
Lachnospiraceae NK4A136 group (B)
Mucor (E)
Muribaculaceae (B)
Bacteroides (B)
Muribaculaceae (B)
Lachnospiraceae (B)
Debaryomyces (E)
Lachnospiraceae NK4A136 group (B)
Lachnospiraceae NK4A136 group (B)
Rikenellaceae RC9 gut group (B)
Enterorhabdus (B)
Lachnospiraceae NK4A136 group (B)
Desmococcus olivaceus (E)
Prevotellaceae UCG-001 (B)
Lachnospiraceae FCS020 group (B)
Magnoliophyta Triticum aestivum (E)
Lachnospiraceae NK4A136 group (B)

**D**

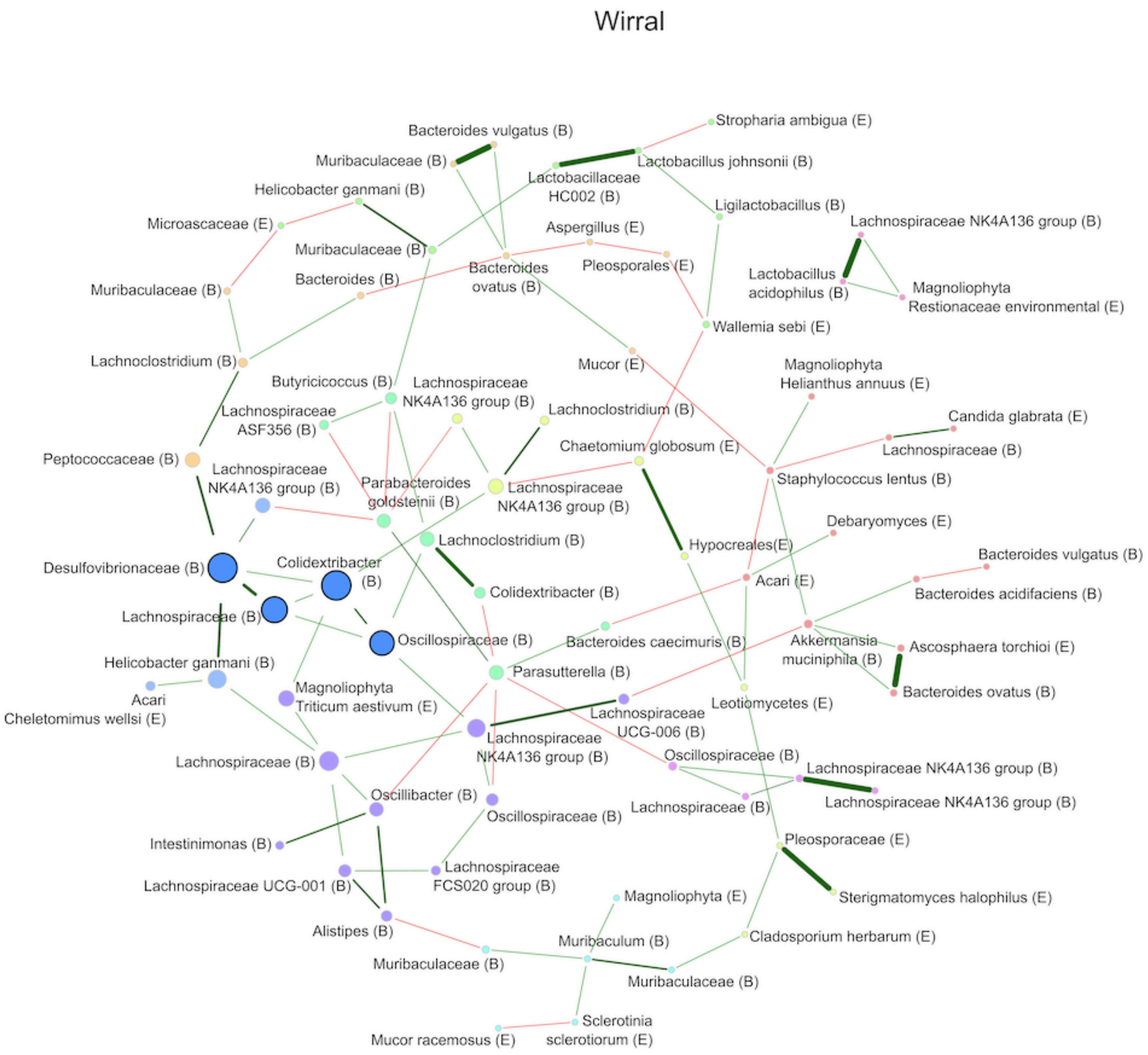
Wirral
Bacteroides vulgatus (B)
Stropharia ambigua (E)
Muribaculaceae (B)
Lactobacillus johnsonii (B)
Lactobacillaceae HC002 (B)
Helicobacter ganmani (B)
Ligilactobacillus (B)
Microascaceae (E)
Aspergillus (E)
Lachnospiraceae NK4A136 group (B)
Muribaculaceae (B)
Bacteroides ovatus (B)
Pleosporales (E)
Bacteroides (B)
Lactobacillus acidophilus (B)
Magnoliophyta Restionaceae environmental (E)
Muribaculaceae (B)
Wallemia sebi (E)
Lachnoclostridium (B)
Mucor (E)
Magnoliophyta Helianthus annuus (E)
Butyricicoccus (B)
Lachnospiraceae NK4A136 group (B)
Lachnoclostridium (B)
Candida glabrata (E)
Lachnospiraceae ASF356 (B)
Chaetomium globosum (E)
Lachnospiraceae (B)
Peptococcaceae (B)
Lachnospiraceae NK4A136 group (B)
Parabacteroides goldsteinii (B)
Lachnospiraceae NK4A136 group (B)
Staphylococcus lentus (B)
Debaryomyces (E)
Lachnoclostridium (B)
Hypocreales(E)
Bacteroides vulgatus (B)
Desulfovibrionaceae (B)
Colidextribacter (B)
Acari (E)
Colidextribacter (B)
Bacteroides acidifaciens (B)
Lachnospiraceae (B)
Oscillospiraceae (B)
Bacteroides caecimuris (B)
Akkermansia muciniphila (B)
Ascosphaera torchioi (E)
Helicobacter ganmani (B)
Parasutterella (B)
Acari Cheletomimus wellsi (E)
Magnoliophyta Triticum aestivum (E)
Bacteroides ovatus (B)
Lachnospiraceae UCG-006 (B)
Leotiomycetes (E)
Lachnospiraceae NK4A136 group (B)
Lachnospiraceae (B)
Oscillospiraceae (B)
Lachnospiraceae NK4A136 group (B)
Lachnospiraceae NK4A136 group (B)
Oscillibacter (B)
Lachnospiraceae (B)
Oscillospiraceae (B)
Intestinimonas (B)
Pleosporaceae (E)
Lachnospiraceae FCS020 group (B)
Lachnospiraceae UCG-001 (B)
Magnoliophyta (E)
Sterigmatomyces halophilus (E)
Alistipes (B)
Muribaculum (B)
Cladosporium herbarum (E)
Muribaculaceae (B)
Muribaculaceae (B)
Sclerotinia sclerotiorum (E)
Mucor racemosus (E)

**Supplementary Information 21.** Example FACS plots of processed faecal samples from (A) a wild mouse, (B) control IgMi mice that do not produce or secrete IgA, and (C) a control of wild mouse faeces without using SYBR Green or the PE-conjugated anti-mouse IgA in the staining protocol. For each, the panels left to right are forward scatter (FSC-A) *vs*. side scatter (SSC); FITC (for SYBR Green) *vs*. FSC-A; PE (for PE-conjugated anti-mouse IgA) *vs*. FSC-A, with the proportional abundance in each category shown in the bottom panel. IgA[hi] and IgA[-] were defined from dot plots of pair-matched negative controls (no IgA staining, as panel C) from each individual, where maximum tolerance to false positives was <0.1%. The IgA[hi] showed a MFI of 11772 (arbitrary unit), sufficiently separated from the IgA[-] (MFI = -829).

**A**

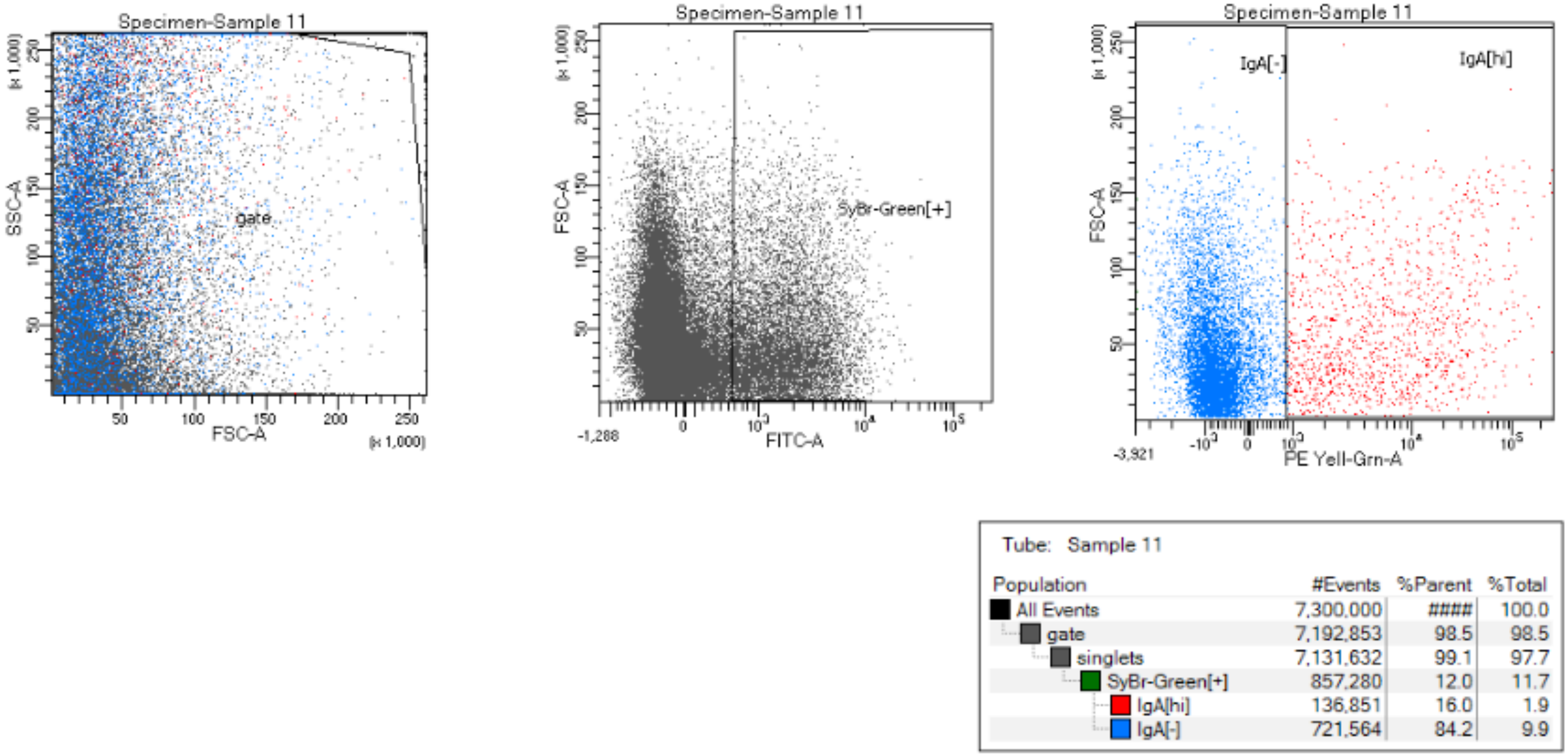

**B**

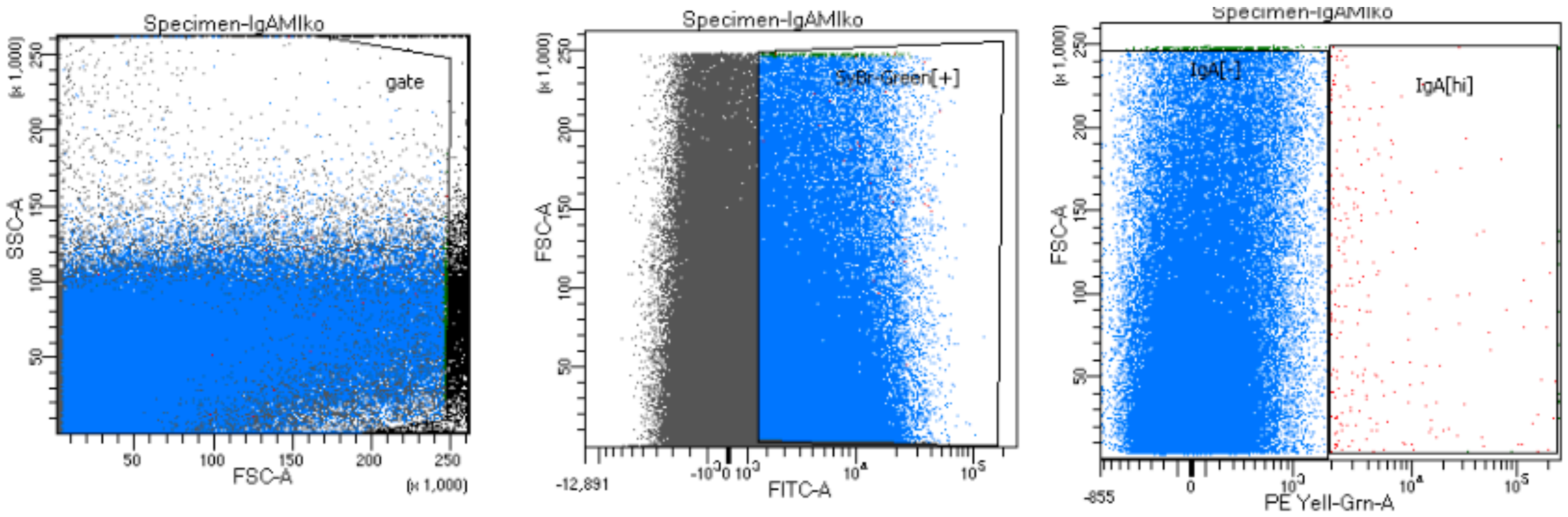

Tube: IgAMIko

| Population | #Events | %Parent | %Total |
|---|---|---|---|
| All Events | 1,313,199 | #### | 100.0 |
| gate | 1,236,058 | 94.1 | 94.1 |
| singlets | 1,231,714 | 99.6 | 93.8 |
| SyBr-Green[+] | 124,603 | 10.1 | 9.5 |
| IgA[hi] | 228 | 0.2 | 0.0 |
| IgA[-] | 124,118 | 99.6 | 9.5 |

**C**

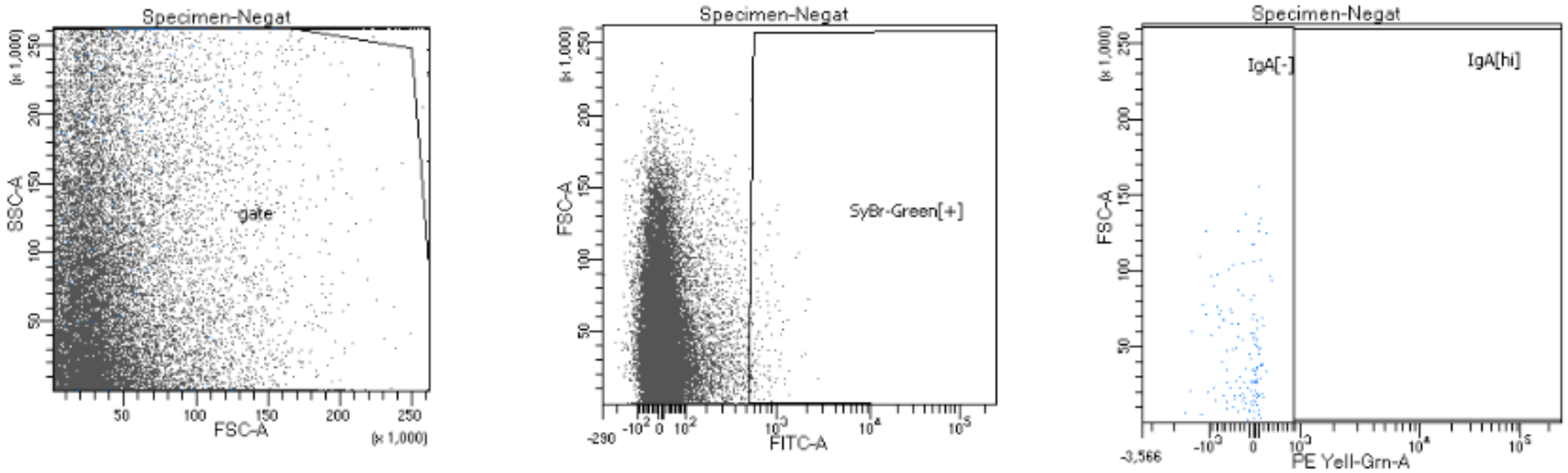

Tube: Negat

| Population | #Events | %Parent | %Total |
|---|---|---|---|
| All Events | 7,300,000 | #### | 100.0 |
| gate | 7,268,048 | 99.6 | 99.6 |
| singlets | 7,182,936 | 98.8 | 98.4 |
| SyBr-Green[+] | 13,717 | 0.2 | 0.2 |
| IgA[hi] | 11 | 0.1 | 0.0 |
| IgA[-] | 13,705 | 99.9 | 0.2 |
